# LAVA: An integrated framework for local genetic correlation analysis

**DOI:** 10.1101/2020.12.31.424652

**Authors:** J. Werme, S. van der Sluis, D. Posthuma, C. A. de Leeuw

## Abstract

Genetic correlation (*r*_g_) analysis is commonly used to identify traits that may have a shared genetic basis. Traditionally, *r*_g_ is studied on a global scale, considering only the average of the shared signal across the genome; though this approach may fail to detect scenarios where the *r*_g_ is confined to particular genomic regions, or show opposing directions at different loci. Tools dedicated to local *r*_g_ analysis have started to emerge, but are currently restricted to analysis of two phenotypes. For this reason, we have developed LAVA, an integrated framework for local *r*_g_ analysis which, in addition to testing the standard bivariate local *r*_g_’s between two traits, can evaluate the local heritability for all traits of interest, and analyse conditional genetic relations between several traits using partial correlation or multiple regression. Applied to 20 behavioural and health phenotypes, we show considerable heterogeneity in the bivariate local *r*_g_’s across the genome, which is often masked by the global *r*_g_ patterns, and demonstrate how our conditional approaches can elucidate more complex, multivariate genetic relations between traits.

## Introduction

Results from just over a decade of genome-wide association studies (GWAS) have demonstrated that statistical pleiotropy across the genome is ubiquitous, meaning that particular genetic variants, genes, or genomic regions often show association with more than one trait^1–3^. Pleiotropy is valuable to study for a number of reasons, as it could elucidate biological pathways that are shared between traits^4–6^, help generate hypotheses about the functional significance of GWAS results^7–9^, and improve our understanding of the aetiology and overlap between complex traits and diseases^1,10,11^.

Pleiotropy on the single variant level has traditionally been studied using colocalization methods, which typically employ a Bayesian analysis framework with the aim of detecting true, causally shared genetic effects^7,8,11,12^; but there now exists a wide range of different cross-trait genetic association methods aimed at elucidating pleiotropy^5,9,13–15^. Given the notion that extensive pleiotropy may result in a genome-wide correlation between the genetic association signals, genetic correlation analysis has been frequently employed to identify traits for which there could be widespread pleiotropy across the genome, and this type of analysis has become a standard follow-up analysis to genome-wide association studies (GWAS)^13,16–18^. Notably, an observed genetic correlation (*r*_g_) does not guarantee the presence of true, causal pleiotropy, as strong linkage disequilibrium (LD) between different causal SNPs could also give rise to a correlation between genetic signals^17^, but genetic correlation analysis nonetheless facilitates prioritisation of scenarios where pleiotropy is likely.

While pleiotropy is typically discussed on a local level, such as single SNPs or genes, *r*_g_ is traditionally studied on a global, genome-wide scale. Since a global *r*_g_ merely represents an average of the shared association signals across the genome, local *r*_g_’s in opposing directions could result in a low or completely non-significant global *r*_g_, and strong local *r*_g_’s in the absence of any global relationship may go undetected^17,19^. In addition, global *r*_g_’s offer limited insight into the biological mechanisms that are shared between phenotypes, as the exact source of any detected genetic relation remains unidentified. To overcome these limitations, some have employed strategies such as partitioning the *r*_g_ by annotation (e.g. GNOVA^20^), or restricted testing only to variants that are assumed to be associated (MiXER^21^).

Due to often high levels of LD between nearby SNPs, global *r*_g_ methods cannot easily be translated to a local scale; but methods aimed at estimating local *r*_g_ have also started to emerge (Rho-Hess^19^, SUPERGNOVA^22^, LOGOdetect^23^). To our knowledge, however, no existing tool currently offers the opportunity to model the local genetic relations between more than two phenotypes simultaneously. To address this, we have developed LAVA (<u>**L**</u>ocal <u>**A**</u>nalysis of [co]<u>**V**</u>ariant <u>**A**</u>nnotation), a flexible and user-friendly tool aimed at detecting regions of shared genetic association signal between any number of phenotypes. LAVA can analyse binary as well as continuous phenotypes with varying degrees of sample overlap, and in addition to evaluating standard bivariate local *r*_g_’s, it can test the local joint univariate association signal (i.e. the local *h*^2^) for the traits in question, which can be used to filter out non-associated loci that may yield unstable *r*_g_ estimates, and analyse the genetic relations between several traits simultaneously. Local genetic association analysis of multiple traits can be performed either via partial correlation or multiple linear regression, allowing for complex, multivariate genetic relationships to be examined in more detail than is currently possible with standard bivariate approaches.

In this paper, we demonstrate the features of LAVA through application to real data, and validate its properties and robustness using simulation. We first describe the details of the method and the various analysis options, and then apply LAVA to 20 behavioural and health-related traits. We examine the heterogeneity of genetic relations between these phenotypes, and then zoom in on the major histocompatibility complex (MHC) on chromosome 6, to examine the variability of the conditional local relationships between selected health phenotypes in this LD-dense region.

## RESULTS

### Input processing and estimation of local genetic signal

For any genomic region of interest, consider a centred continuous phenotype vector *Y_p_* (for phenotype *p*) with sample size *N*, and a standardised genotype matrix *X* containing K_’()_ SNPs. We can model the relation between a phenotype and all SNPs in this region using a multiple linear regression model of the form *Y*_*p*_ = *Xα_p_* + ∈_*p*_, where α_*p*_ represents the vector of joint SNP effects (which account for the LD between SNPs) and ∈_*p*_ the vector of residuals which are normally distributed with variance 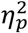

Given that the standard least squares estimate of α_*p*_ is of the form 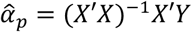, if we denote the local SNP LD matrix as *S* = cor*(X)*and the vector of estimated marginal SNP effects as 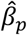 (which do not account for LD), we can express 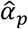 as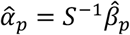. Then, after obtaining 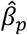 from GWAS summary statistics for *Y_p_*, by using a reference genotype data set (e.g. 1,000 Genomes^24^) from a population with a matching ancestry/LD structure to compute *S*, we can estimate the joint SNP effects α_*p*_ (effectively removing the correlation between SNPs effects that is due to LD), without the need for any individual level data. To ensure that the direction of effect is consistent across traits (which is crucial for preventing false positives; see **Suppl. Fig. 1**), LAVA aligns the summary statistics to the reference data before computing 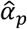

Once we have obtained 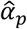, we can estimate the residual variance 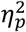, and hence also the phenotypic variance explained by the SNPs in the locus (i.e. the univariate local genetic signal, or local *h*^2^). To determine whether the local *h*^2^ is significant, we test the explained phenotypic variance using an F-test (see Methods). We recommend using this test to filter out non-associated loci prior to any *r*_g_ analysis, since *r*_g_’s will not be interpretable or reliable for phenotypes that do not show any local genetic signal.

Note that the above regression formulation concerns continuous phenotypes; for binary phenotypes we employ a largely similar strategy, reconstructing a multiple logistic regression model for the locus based on the marginal SNP effects and using a *X*^*2*^-test to test the joint association of SNPs with the phenotype (see Methods for more detail).

### Estimation of bivariate local genetic correlations

To determine the bivariate local genetic correlation: for any region and set of *P* traits, we define the local genetic component matrix *G = X*α, where *X* represents the standardised genotypes at that locus and α the K_snp_ by *P* matrix of joint effects on each trait (as outlined above). We denote the realised covariance matrix of *G* as Ω, such that each diagonal element 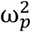 represents the local genetic variance of *G*_p_ for phenotype *p*, and each off-diagonal element *ω*_pq_ the local genetic covariance of *G*_p_ and *G*_q_ (for phenotypes *p* and *q*). Thus, for two phenotypes *p* and *q*:

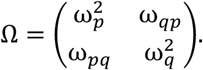

From this Ω, the local genetic correlation can be directly computed as 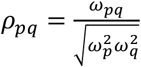. Since genotype and phenotype data is all standardised, the square of the estimated local genetic correlation 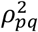, can be interpreted as the proportion of variance in the local genetic component *G* that is explained by *G*_q_ (and vice versa). Since *G* is unobserved, Ω cannot be computed directly, and we therefore estimate it using the Method of Moments^25^. Once estimated, we compute *ρ*_*pq*_ as shown above, and generate simulation-based based *p*-values to evaluate its significance (see Methods).

As shown in **Suppl. Figs. 2-4**, this approach produces unbiased parameter estimates with well contained type 1 error rates for both binary and continuous phenotypes, and a wide range of locus sizes. See also the **Suppl. Note 1** for a comparison of the local *r*_g_ estimation employed in LAVA, to that of Rho-Hess and SUPERGNOVA.

Since any potential sample overlap between summary statistics sets can result in an upward bias in the estimated correlation, known or estimated sample overlap (obtained e.g. via bivariate LDSC^13^) should be provided to LAVA. Any shared variance that is due to sample overlap will be modelled as a residual covariance, effectively removing such bias (see Methods; **Suppl. Fig. 5**).

### Estimation of multivariate local genetic relations

Multivariate local genetic analysis can be used to obtain the conditional genetic associations, using several traits simultaneously. This has been implemented in two forms: partial correlation, which models the local *r*_g_ between two phenotypes while controlling for their *r*_g_’s with one or more other phenotypes, and multiple regression, which can model local genetic signal of an outcome phenotype using a set of predictor phenotypes simultaneously.

The partial genetic correlation between the phenotypes *p* and q, conditioned on their local *r*_g_’s with some other phenotypes(s) Z, is denoted ρ_*pq|z*_. This ρ_*pq|z*_ can be computed directly from Ω (see Methods), and indicates how much of the initial correlation ρ_*pq*_ remains once the *r*_g_’s with the phenotypes(s) in *Z* are accounted for.

In contrast, the multiple regression models the genetic signal for a single outcome phenotype Y using the genetic signal for one or more predictor phenotypes *X*. We formulate this as *G*_Y_ *= G*_X_γ + ε for standardised genetic components *G*_X_ and *G*_Y_, such that γ reflects the vector of standardised regression coefficients and ε the residuals, with residual variance τ^*2*^ (all computed directly from Ω; see Methods). Here, the γ’s indicate how much the genetic component for each individual predictor in *X* contributes to the genetic component of *Y* (conditioned on the other predictors) and τ^*2*^ the proportion of local heritability of *Y* that is independent of *X*. From this τ^*2*^, we can then compute the model *r*^*2*^ as *r*^*2*^ *=* 1 – *τ*^*2*^, which tells us how much of the local heritability for *Y* can be explained by the genetic components of all predictor phenotypes jointly. (95% confidence intervals are computed for all individual γ’s, as well as the total model *r*^*2*^; Methods). For a more in-depth overview of the differences and similarities between partial correlation and multiple regression, see **Suppl. Note 2**.

As can be seen in **Suppl. Figs. 6-9**, our two multivariate local genetic association approaches provide overall unbiased estimates and controlled type 1 error rates. The only exceptions were a few instances of the partial correlation for binary phenotypes where we saw a minor median-bias, although no mean-bias, for some null simulations with very low univariate signal (although type 1 error rates for these were nevertheless controlled). Additionally, there was a very slight type 1 error rate inflation at a local odds ratio of 1.5, which had an error rate of .06 for an alpha level of .05 (**Suppl. Fig. 9**), though we note that this represents a rather extreme level of local heritability for complex, non-mendelian phenotypes.

### Bivariate local genetic correlation analyses reveal extensive overlap of local genetic association signals between traits

To demonstrate our method, we applied LAVA to 20 health related and behavioural traits (see **Table 1**), testing the pairwise local genetic correlations within 2,495 genomic loci (genome-wide), followed by conditional local genetic analyses for a subset of strongly intercorrelated phenotypes. In order to partition the genome, we developed an algorithm that sections the chromosomes into approximately equal sized (∼1Mb) semi-independent blocks, by recursively splitting the chromosomes into smaller regions while minimising the LD between them (see Methods – Genome partitioning for details).

As our summary statistics were based on European samples, we utilised the individuals of European ancestry from the 1000 Genomes (phase 3)^24^ as a genotype reference (both for the definition of genomic regions and for all LAVA analyses). Sample overlap was estimated using the intercepts from bivariate LDSC (see Methods).

Given that the detection of valid and interpretable local *r*_g_’s requires the presence of sufficient local genetic signal, we used the univariate test as a filtering step for the bivariate local *r*_g_ analyses. Since the power to detect a significant local heritability depends on the power of the original GWAS, this step could potentially lead to the exclusion of some relevant loci, particularly for phenotypes with a small sample size. Though similar to a lack of genetic signal, such scenarios would likely also produce unstable *r*_g_ estimates, and we therefore tested only the local correlations for any pairs of traits which both exhibited univariate local genetic signal at *p* < .05 / 2,495. This resulted in a total of 21,374 bivariate tests across all trait pairs, spanning 1,919 unique loci.

With a Bonferroni corrected *p*-value threshold of 2.34e-6 (.05 / 21,374), we detected a total of 546 significant bivariate local *r*_g_’s across 234 loci, of which 81 loci were associated with more than one trait pair. For 193 of these correlations, the 95% confidence intervals (CI’s) for the explained variance included 1, which is consistent with the scenario that the genetic signal of those pairs of phenotypes in these loci is completely shared (**Fig. 1**).

**Figure 1:**
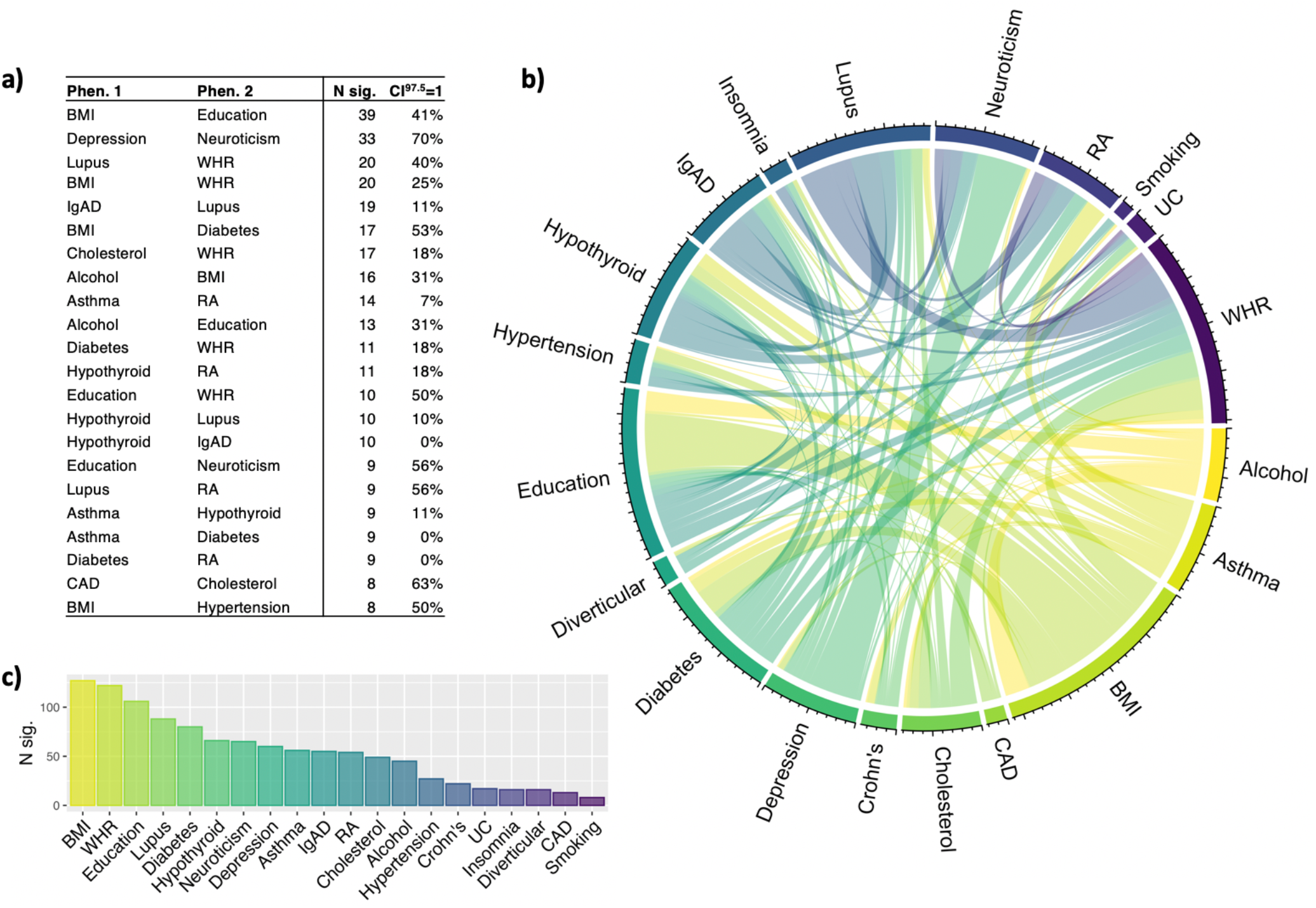
Overview of the number of significant bivariate local r_g_’s between all 20 trait pairs. The table (a) shows the exact number of significant bivariate local r_g_’s detected between top phenotype pairs (N. sig.), together with proportion of significant loci for which the 95% confidence interval included 1 (CI^97.5^=1). The chord diagram (b) illustrates the number of significant bivariate local genetic correlations between all phenotype pairs, while the bar plot (c) shows the total number of significant bivariate correlations detected per phenotype.

The trait pairs exhibiting the greatest number of significant local *r*_g_’s were BMI and educational attainment (39), which also had with the largest sample sizes (see **Table 1**), followed by depression and neuroticism (33), and BMI and waist-hip ratio (20). As can be seen in **Fig. 1**, several conceptually related phenotypes tended to show a large number of significant *r*_g_’s, with depression and neuroticism having the highest proportion of regions within which the CI’s included 1.

**Table 1.** Overview of the 20 phenotypes included in this study

| **Phenotype** | **Short ID** | **N** | **N cases** | **N controls** | **Global $h^2$ (SE)** | **Atlas ID** | **Original study** |
| --- | --- | --- | --- | --- | --- | --- | --- |
| **High cholesterol** | *Cholesterol* | 289,307 | 46,932 | 242,375 | 5% (.5) | 3608 | *Watanabe, K. et al. (2019)<sup>2</sup>* |
| **Coronary artery disease (incl. angina)** | *CAD* | 86,995 | 22,233 | 64,762 | 4% (.3) | 3925 | *Schunkert, H. et al. (2011)<sup>27</sup>* |
| **Hypertension** | *Hypertension* | 244,890 | 71,332 | 173,558 | 10% (.5) | 3691 | *Watanabe, K. et al. (2019)<sup>2</sup>* |
| **Crohn's disease** | *Crohn's* | 20,883 | 5,956 | 14,927 | 48% (5.5) | 68 | *Liu J. Z. et al. (2015)<sup>28</sup>* |
| **Ulcerative colitis** | *UC* | 27,432 | 6,968 | 20,464 | 25% (3.3) | 69 | *Liu J. Z. et al. (2015)<sup>28</sup>* |
| **Diverticular disease** | *Diverticular* | 451,099 | 31,964 | 419,135 | 3% (.2) | 4311 | *Schafmayer, C. et al. (2019)<sup>29</sup>* |
| **Body mass index** | *BMI* | 379,831 | - | - | 22% (.7) | 3445 | *Watanabe, K. et al. (2019)<sup>2</sup>* |
| **Waist-hip ratio (adj. for BMI)** | *WHR* | 694,649 | - | - | 11% (.5) | 4080 | *Pulit, S. L. et al. (2019)<sup>30</sup>* |
| **Diabetes** | *Diabetes* | 385,420 | 18,483 | 366,937 | 4% (.3) | 3328 | *Watanabe, K. et al. (2019)<sup>2</sup>* |
| **Asthma** | *Asthma* | 385,822 | 44,301 | 341,521 | 5% (.5) | 3552 | *Watanabe, K. et al. (2019)<sup>2</sup>* |
| **Hypothyroidism** | *Hypothyroid* | 244,890 | 13,043 | 231,847 | 4% (.6) | 3685 | *Watanabe, K. et al. (2019)<sup>2</sup>* |
| **Rheumatoid arthritis** | *RA* | 58,284 | 14,361 | 43,923 | 21% (7.4) | 1203 | *Okada, Y. et al. (2013)<sup>31</sup>* |
| **Lupus** | *Lupus* | 14,267 | 5,201 | 9,066 | 84% (21.1) | 4018 | *Julià, A. et al. (2018)<sup>32</sup>* |
| **Immunoglobulin A deficiency** | *IgAD* | 6,487 | 1,635 | 4,852 | 78% (42.3) | 2037 | *Bronson, P. G. et al. (2016)<sup>33</sup>* |
| **Depression** | *Depression* | 500,199 | 170,756 | 329,443 | 6% (.2) | 4293 | *Howard, D. M. et al. (2019)<sup>34</sup>* |
| **Neuroticism** | *Neuroticism* | 380,506 | - | - | 10% (.4) | 3990 | *Nagel, M. et al. (2018)<sup>35</sup>* |
| **Insomnia** | *Insomnia* | 386,078 | - | - | 6% (.2) | 3232 | *Watanabe, K. et al. (2019)<sup>2</sup>* |
| **Educational attainment** | *Education* | 766,345 | - | - | 10% (.3) | 4066 | *Lee, J. et al. (2018)<sup>36</sup>* |
| **Alcohol intake frequency** | *Alcohol* | 386,082 | - | - | 7% (.3) | 3261 | *Watanabe, K. et al. (2019)<sup>2</sup>* |
| **Current tobacco smoking** | *Smoking* | 386,150 | - | - | 4% (.2) | 3235 | *Watanabe, K. et al. (2019)<sup>2</sup>* |

Summary statistics have been downloaded from the GWAS Atlas<sup>2</sup>, the relevant page for individual summary statistics can be accessed directly at: https://atlas.ctglab.nl/taitDB/[Atlas-ID]. SNP heritabilities ( $h^2$ ) were obtained using LDSC<sup>37</sup>.

Given the number of immune phenotypes analysed here, we chose to retain the major histocompatibility complex (MHC; chr6:26-34Mb, 21 loci) in our analyses as this locus is highly relevant to the aetiology of these phenotypes. Of the 546 significant local *r*_g_’s, 229 were found within these MHC loci (particularly for immune phenotypes), consistent with the notion that there is extensive pleiotropy within this region^2,26^.

### Local genetic correlation analysis more accurately captures the heterogeneous genetic relationships between phenotypes

For all trait pairs, we examined the strength and direction of effect of the local *r*_g_’s by taking the average of the observed correlation coefficients across tested loci. As shown in **Fig. 2**, consistently positive *r*_g_’s with multiple significant loci were observed for many phenotypes (e.g. neuroticism & depression, cholesterol & CAD, BMI & diabetes, Crohn’s & UC), for which we also saw concordance with the observed pattern of global *r*_*g*_’s, as obtained via bivariate LDSC^13^ (**Fig. 2**).

**Figure 2:**
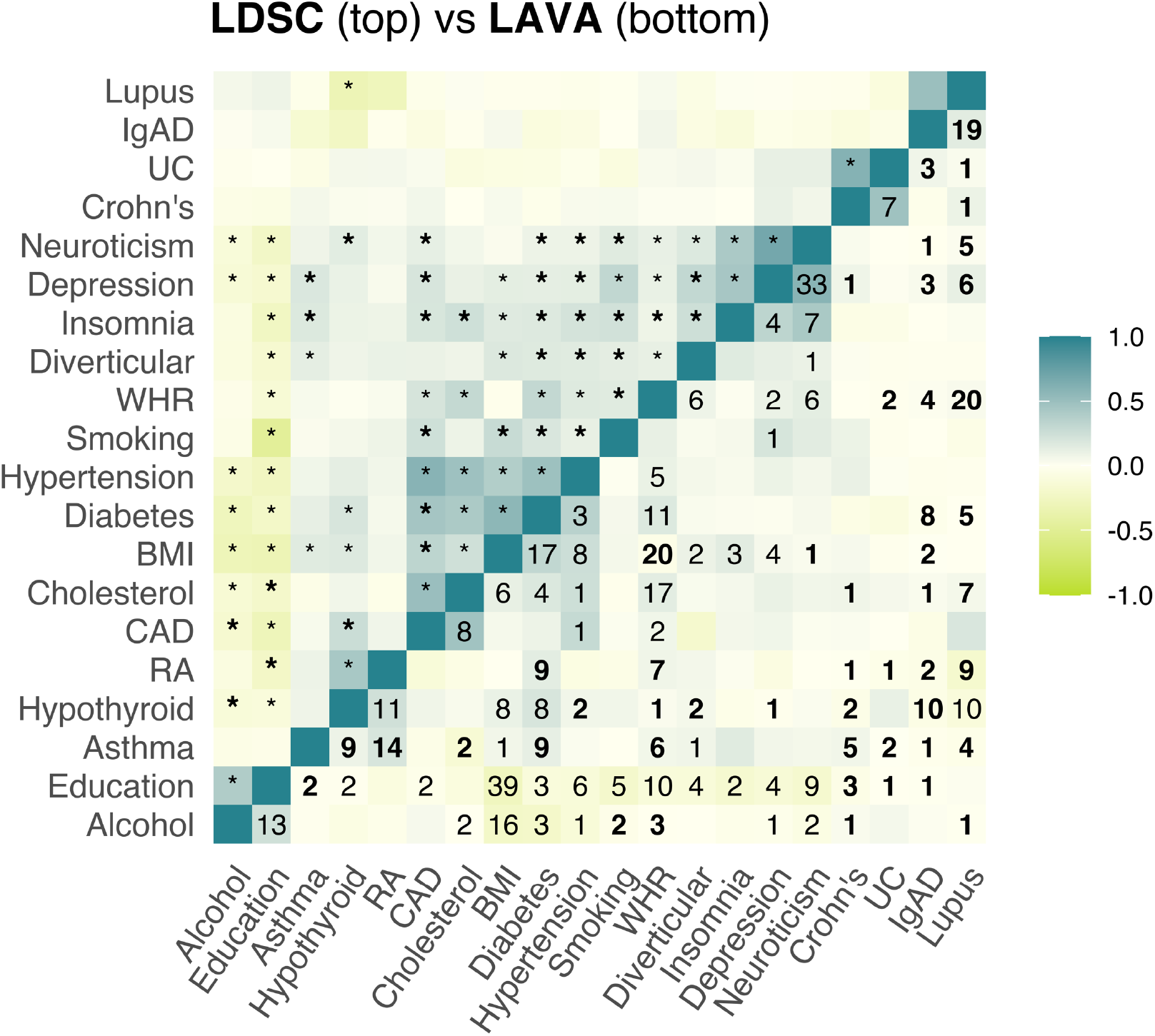
Comparison between the global genetic correlation estimated using LDSC (top) and the mean local genetic correlation from LAVA across tested loci (bottom). The “*” indicates a significant global genetic correlation estimated by LDSC (p < .05 / 190 = 2.63e-4), while the numbers indicate how many significant local r_g_’s were detected with LAVA (p < .05 / 20,630 = 2.42e-6). Bold asterisk means a significant global r_g_ was detected by LDSC, without any evidence for a significant local r_g_, while bold numbers indicate that at least one local r_g_ was detected by LAVA, despite no significant global r_g_. While LDSC excludes a small region within the MHC (chr6:30-31Mb), this region concerns only three of the loci analysed with LAVA (locus IDs: 955-957), within which we detected 14 significant r_g_’s between 10 traits: all of which had significant r_g_’s elsewhere.

However, there were also several trait pairs with a global *r*_g_ close to 0 which nonetheless exhibited significant local genetic correlations (e.g. BMI & WHR, BMI & neuroticism, asthma & Crohn’s, asthma & UC, alcohol & WHR), supporting the notion that global *r*_g_’s fail to capture the complexity and heterogeneity in the genetic overlap between many traits. As expected, these traits tended to exhibit local *r*_g_’s in opposite directions, and/or within a limited number of loci. Although even for pairs like alcohol intake frequency and BMI, which showed a consistent pattern of negative local *r*_g_’s, and a highly significant negative global *r*_g_ (*r* = -.3, *p* = 8.85e-42), two significant local *r*_g_’s in a positive direction were still found. Similar patterns were observed for several other phenotypes, such as BMI and cholesterol, diabetes and cholesterol, and alcohol and neuroticism (for an overview of the total number of positive versus negative local genetic correlations detected per phenotype pair, see **Suppl. Fig. 10**).

### Bivariate local genetic correlations implicate potential pleiotropy hotspots

From the bivariate analyses, we identified a total of 81 regions within which significant *r*_g_’s were found between multiple trait pairs. Most of these were located within the MHC (chr6:26-34Mb), a region within which extensive pleiotropy has been noted previously^2,26^. Within MHC hotspots, immune related phenotypes were among the most frequently intercorrelated (with lupus displaying the greatest number of significant genetic correlations of all; **Fig. 3**), consistent with the known role of the MHC in immune function^38,39^.

**Figure 3:**
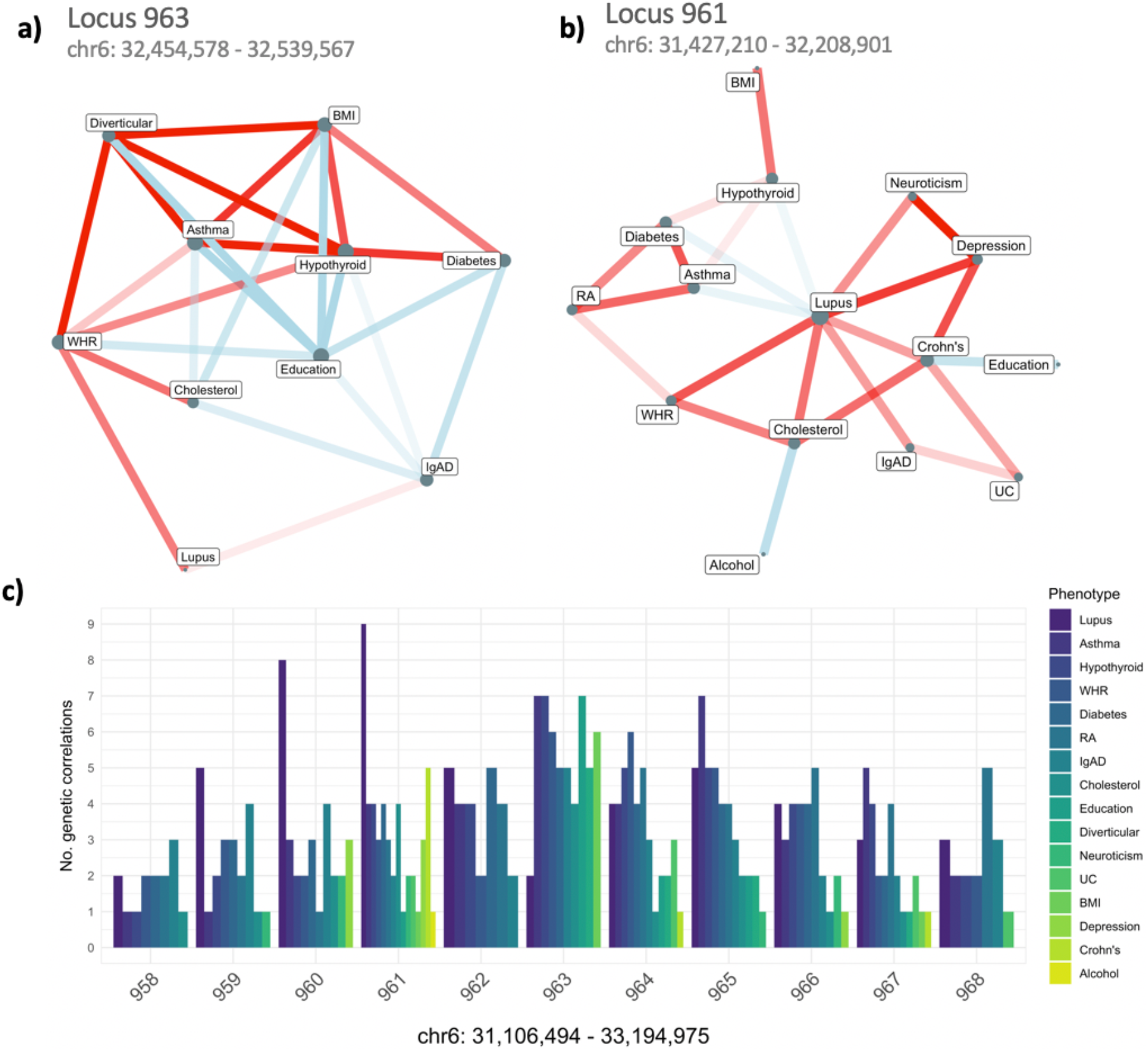
Dense bivariate local genetic correlations within the MHC. The network plot (a) shows the bivariate local r_g_’s within the two biggest MHC hotspots (locus 963, chr6:32,454,578-32,539,567 & locus 961, chr6:31,427,210-32,208,901). Colour indicates direction of effect (red = positive, blue = negative) and opacity the strength, while node size reflects the number of significant local r_g_’s for each phenotype. The barplot (b) shows how many significant bivariate r_g_’s were detected for each phenotype on the Y-axis, within the 10 adjacent loci of the biggest hotspot on the X axis (chr6:31,106,494-33,194,975). The phenotypes have been ordered according to total number of significant r_g_’s across these loci, indicating that lupus, asthma, hypothyroidism, WHR, and diabetes were the most interconnected within this region.

The largest hotspot, i.e. the locus with the greatest number of genetic correlations, was locus 963 (chr6:32,454,578-32,539,567, within the MHC), where a total of 27 significant correlations between 10 different phenotypes were detected (**Fig. 3a; Suppl. Table 1**). This locus contains a single protein coding gene, HLA-DRB5, which has been linked with several of the associated phenotypes previously (e.g. asthma^40^, diabetes^6^, WHR^41^, lupus^42^). The second largest hotspot, locus 961 (chr6:31,427,210-32,208,901; also within the MHC), was the most diverse, showing a total of 24 significant genetic correlations for 15 different traits. Here, lupus was situated as a hub phenotype, showing significant correlations with most other phenotypes (**Fig. 3b**), a pattern that was observed across a few other MHC loci as well (**Suppl. Tables 2**,**6**,**9**,**13**). Notably, both of these loci were contained within a region identified as the top pleiotropic locus in a recent large scale investigation of pleiotropy by Watanabe et al. (2019)^2^, across a total of 558 traits.

Although the MHC is a region known for its complex LD structure^43^, we nonetheless observed clustering of conceptually related traits within these loci (e.g. cholesterol & WHR vs Crohn’s & UC, depression & neuroticism see **Suppl. Tables 1-11, 13, 15-18, 21, 25-27**), and saw instances of substantial local heritability, without necessarily the presence of any local genetic correlation (e.g. IgAD and RA in loci 958-960: univariate *p*’s < 1e-118, bivariate, bivariate *p*’s > .05). This suggests that shared and nonshared genetic signal might be distinguishable even in regions with notoriously strong LD; though, it should be noted that in scenarios where separate causal SNPs are in perfect LD, true pleiotropy will be inseparable from confounding, and such instances might be more common within LD-dense regions like the MHC.

Outside the MHC, the two largest hotspots had 8 significant *r*_g_’s each. The first on chromosome 11 (112,755,447-113,889,019), with phenotypes depression, neuroticism, alcohol intake, educational attainment, and WHR (locus ID 1719; **Suppl. Table 12**), and the second on chromosome 3 (47,588,462-50,387,742) with educational attainment, insomnia, alcohol intake, BMI, CAD, UC, and Crohn’s (locus ID 464; **Suppl. Table 14**). These hotspots also overlapped with loci identified among the top pleiotropic regions for psychiatric, cognitive, metabolic, and immunological phenotypes previously (see Suppl. Table 4 in Watanabe et al. 2019^2^). In addition, locus 1719 on chromosome 11 contains both *NCAM1* and *DRD2* (among 8 other genes), which have been frequently implicated in behavioural and psychiatric traits (e.g. alcohol dependence^44^, smoking^2^, cannabis use^45^, depression^34^, neuroticism^2,46^, sleep duration^47^, ADHD symptoms^48^), and this locus also overlapped with a hotspot flagged by SUPERGNOVA^22^ within which significant *r*_g_*’s* were identified for autism, bipolar disorder, depression, cognitive performance, schizophrenia and smoking initiation^22^, suggesting this region might be a key regulator of brain related phenotypes.

Finally, we also identified two loci that were specific to RA, hypothyroidism, and lupus: the first on chromosome 2 (191,051,955-193,033,982; locus ID 374) and the second on chromosome 1 (113,418,038-114,664,387; locus ID 100) (see **Suppl. Tables 40 & 42**). In both cases, positive *r*_g_*’s* were observed between hypothyroidism and RA, both of which showed negative *r*_g_*’s* with lupus.

For a complete overview of all hotspots, including the relevant statistics, associated genes, and network plots, see **Suppl. >Tables 1-81**.

### Genetics of asthma partially explain localgenetic correlations between other health-related phenotypes

To demonstrate the application of our partial local *r*_g_ approach – which can model the local *r*_g_ between two phenotypes of interest conditioned on one or more other phenotypes – we selected a subset of the aforementioned MHC hotspots within which we discovered a sub-cluster of four phenotypes that were consistently interconnected: asthma, hypothyroidism, RA, and diabetes (see network plots in **Suppl. Tables 3-5**,**8**). Given that asthma tended to show consistently strong *r*_g_’s with the other phenotypes within this cluster (see **Fig. 4 & Suppl. Tables 3-5**,**8**), we hypothesized that the shared genetic signal with asthma could account for some of the overlap between RA, diabetes, and hypothyroidism. We therefore computed the partial genetic correlations between these phenotypes, accounting for their local *r*_g_ with asthma.

**Figure 4:**
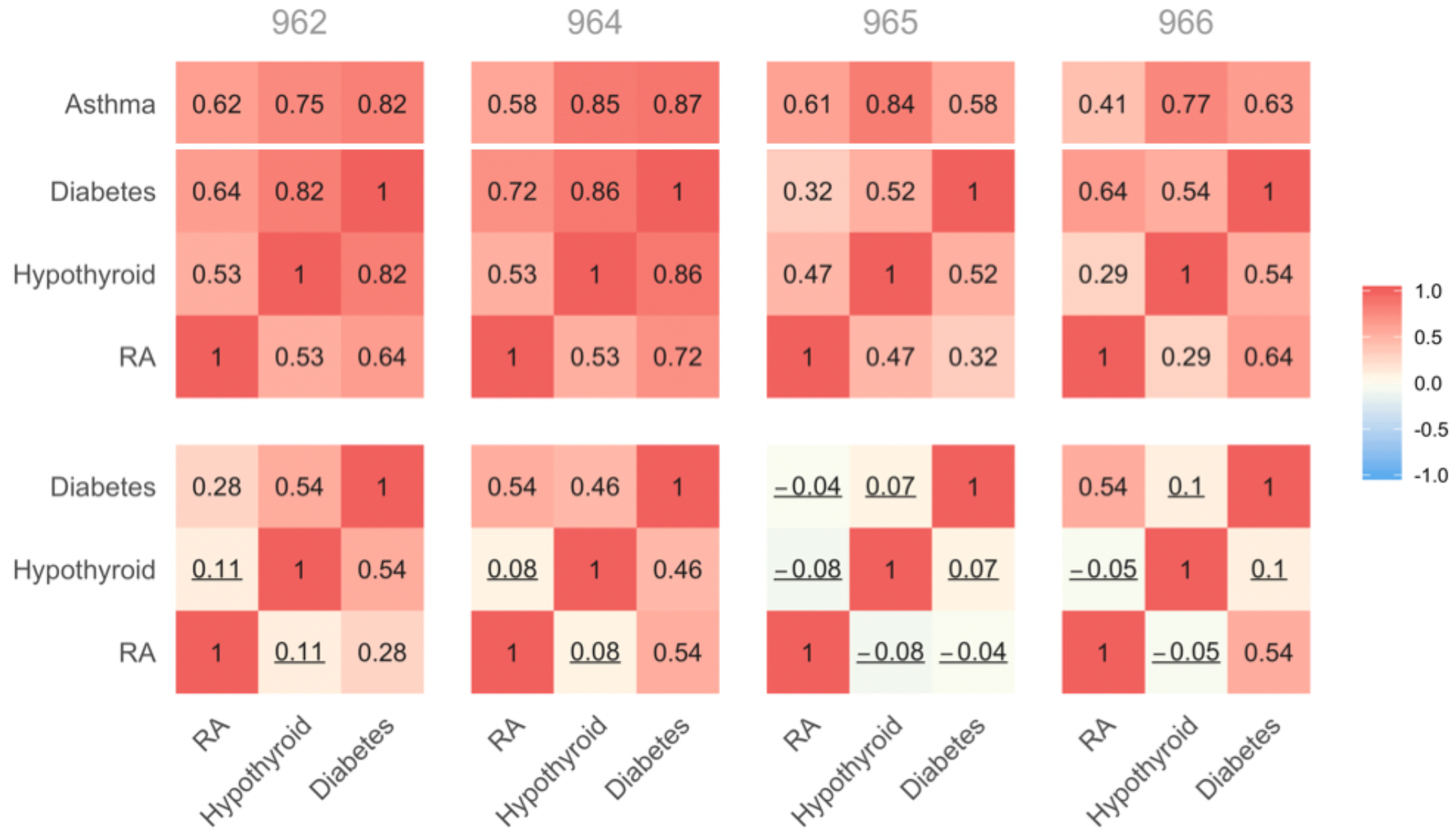
Heatmaps demonstrating the effect of conditioning on asthma within four MHC loci (chr6: 32,208,902–32,682,213). The top heatmaps show the unconditioned bivariate local r_g_, while the bottom heatmaps show the partial local r_g_’s conditioned on asthma. Correlation coefficients with 95% confidence intervals spanning 0 are indicated with an underscore.

**Figure 5:**
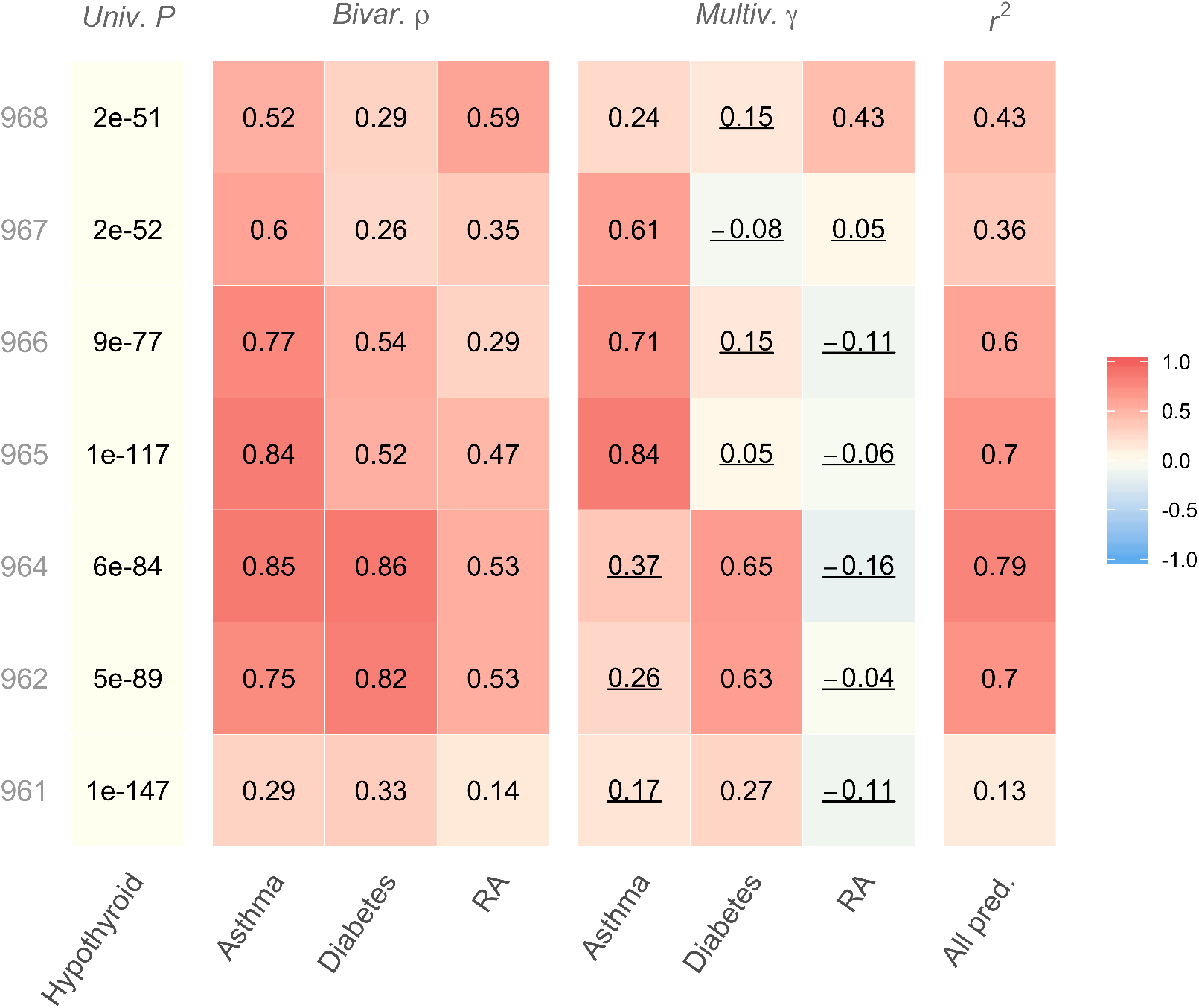
Results for multiple regression model of hypothyroidism on six other genetically correlated traits within MHC loci. The heatmap shows the bivariate genetic correlations (ρ) as well as the standardised coefficients (γ) from the multivariate regression models (together with the multivariate r^2^) within 7 MHC loci (chr6: 31,427,210-33,194,975Mb; Note, there were too few common SNPs in locus 963 to perform the multivariate analysis in that locus). Parameters for which the 95% confidence intervals included 0 are underscored. None of the CI’s for the multivariate r^2^’s included 1, suggesting the presence of hypothyroidism-specific genetic signal in all seven loci.

As shown in **Fig. 4**, conditioning on asthma, in most cases, resulted in a substantial decrease of the *r*_g_’s between hypothyroidism, RA, and diabetes. On several occasions, the 95% CI’s for the partial *r*_g_’s included 0, indicating that they were no longer even nominally significant. This reduction in signal was particularly evident for locus 965 (chr6:32,586,785-32,629,239), in which the partial *r*_g_’s were no greater than .08 after accounting for asthma (and all of the CI’s spanning 0), despite bivariate *r*_g_’s ranging from .32 to .52. This suggests that, in part, genetic variants associated with asthma within these loci might confer a more general susceptibility these phenotypes.

Notably, the degree of change in the local *r*_g_ after conditioning on asthma was locus dependent, indicating variation in the local genetic covariance structure even for adjacent blocks within the MHC.

### Shared genetic aetiology of hypothyroidism within the MHC

To also demonstrate the multiple regression approach – which can model the genetic signal for a single outcome phenotype of interest using the genetic signal for one or more predictor phenotypes – we selected hypothyroidism as the outcome and computed the full joint local genetic relations across a set of relevant MHC loci, using asthma, diabetes, and rheumatism as predictors. With this, we determined the total proportion of variance in the genetic component of hypothyroidism that can be attributed to the genetic signal for these three traits simultaneously.

As seen in **Fig. 5**, there was notable variation in the total multivariate *r*^2^ across adjacent MHC loci, with the proportion of the genetic component of hypothyroidism that could be explained by that of the predictor phenotypes ranging from as little as 13% to as much as 79%. Note, however, that the CI’s for the joint *r*^2^’s did not include 1, suggesting that some proportion of the local heritability for hypothyroidism within these loci is nonetheless independent of the three predictor phenotypes.

There was also substantial variation in the strength of the effects of the three traits on hypothyroidism in the multivariate model, largely mirroring the bivariate correlations. In general, either asthma or diabetes tended to account for most of the shared association signal across loci, with 95% CI’s for the multiple regression coefficients of other traits all spanning 0. The relationship between RA and hypothyroidism was largely accounted for by either of asthma or diabetes in all but locus 968 (chr6:32,897,999-33,194,975) where there was some independent association signal for both asthma and RA.

## DISCUSSION

Global genetic correlation (*r*_g_) analysis is commonly used to identify pairs of traits that have a shared genetic basis, and is a widely popular follow-up to GWAS. The traditional, global approach to *r*_g_ analysis reports only the average *r*_g_ across the genome, and may therefore fail to detect more complex and heterogeneous genetic relationships where the signal might be confined to specific regions or even show opposing association patterns at different loci^13,17,19^.

Here, we presented a novel method, LAVA, which is an integrated statistical framework aimed at testing the local genetic relations within user-defined genomic regions. LAVA handles both continuous and binary phenotypes with varying degrees of sample overlap, and in addition to computing standard bivariate local *r*_g_’s between two phenotypes, LAVA can test the local univariate genetic association signal for each phenotype, and model the conditional local genetic relations between several traits simultaneously using either partial correlation or multiple linear regression.

Applied to 20 different behavioural and health related traits across 2,495 semi-independent regions defined based on LD, we identified a total of 546 significant bivariate local *r*_g_’s across 234 regions. Although the direction of effect for individual pairs was in many cases consistent across loci (particularly for traits showing a strong global *r*_g_), there was substantial variability in the strength of the association across the genome, indicating that the genome-wide *r*_g_ is far from constant. In addition, we identified significant *r*_g_’s in opposing directions for several phenotypes, implying a more complex aetiological relationship than that revealed by a global *r*_g_ analysis. Significant local *r*_g_’s were also observed between several trait pairs whose global correlation was not significant, further emphasizing the value of stratifying *r*_g_ by region.

From the bivariate local *r*_g_ analyses, we identified several regions that harboured significant *r*_g_’s between multiple trait pairs, implicating these regions as potential pleiotropy hotspots. As expected, many of these hotspots were located in the MHC, likely owing to the number of immune- and health related phenotypes included in our example (with the MHC frequently implicated in immune function^38,39^), and the MHC having been flagged as a pleiotropy hotspot in the past^2,26^.

We emphasize that while the aim of an *r*_g_ analysis is to elucidate pleiotropy, the ability to do so is naturally limited by the amount of LD that exists within a region. While this LD structure is accounted for by LAVA, there may be cases where distinct causal scenarios yield identical patterns of SNP associations within a locus, and with the extensive LD that exists within the MHC, there is an increased chance that true pleiotropy may be indistinguishable from confounding here. In spite of this, however, we did observe substantial levels of univariate genetic signal without necessarily the presence of any genetic correlation, with the *r*_g_ patterns reflecting some clustering of conceptually related traits, suggesting that strong genetic signal may be distinguishable from genetic covariance even within LD dense regions such as the MHC. However, experimental evidence will be required to confirm these observations.

Based on the elaborate *r*_g_’s patterns observed within the MHC, we selected a subset of loci within this region to demonstrate how more complex association patterns can be disentangled via our two multivariate models: the partial correlation, which tests the genetic correlation between two phenotypes of interest conditioned on some other phenotype(s), and the multiple linear regression, which models the genetic signal of an outcome phenotype using that of several predictor phenotypes jointly (i.e. conditioned on each other). For a cluster of consistently associated phenotypes – asthma, diabetes, RA, and hypothyroidism – we showed how such models allow us to examine in detail the patterns of mediation and confounding that exist between them; providing further insights into the genetic association between traits beyond what can be achieved using standard bivariate models.

The LAVA analysis framework can be applied to answer a wide array of research questions. It may be used in a more targeted manner to follow up on a smaller subset of loci highlighted through GWAS, identifying regions of shared association with aetiologically informative phenotypes, or in a more agnostic manner, scanning multiple traits across the entire genome (as done in this paper). Approaching the genomic region as the unit of interest, LAVA could be applied to study the function of particular blocks or genes by mapping out patterns of genetic sharing within a locus across the phenome (similar to a PheWAS). This general analysis framework will have implications for our understanding of disease aetiology and genetic heterogeneity as a whole, which can be further aided by integrating summary statistics of molecular phenotypes or endophenotypes (such as gene expression, metabolites, or brain regions), facilitating the functional interpretation of GWAS results by evaluating the local *r*_g_’s with these lower level phenotypes. In this setting, the conditional models could prove particularly useful as they may enable identification of key tissues or regions, offering unique insight into the biological mechanisms that underlie complex traits.

Our method is not without limitations. As already discussed, analytical approaches like LAVA can only pinpoint locations where pleiotropy is likely, but these may be confounded by excessive LD and, ultimately, experimental validation will be required to establish the true nature of any observed genetic overlap. In addition, significant local genetic correlations could be detected from multiple nearby regions, but as LAVA can currently only analyse a single locus at a time, it is unable to condition on the association signal from nearby loci, and it is therefore possible that local genetic correlations are observed in regions adjacent to those harbouring the true association signal. LAVA is also limited by the number of overlapping SNPs within different summary statistics data sets, which could potentially lead to a failure to detect true correlations in scenarios where there are too few shared SNPs between them. Though we endeavour to address these limitations as best possible in future versions of LAVA.

## METHODS

### Model overview and input processing of continuous phenotypes

For any given locus and phenotype *p*, consider a linear regression model of the standardised phenotype vector *Y*_*p*_ on a genotype matrix *X* (containing *K*_snp_ SNPs, also standardised): *Y*_*p*_ = *Xα*_*p*_ + *ϵ*_*p*_, where *α*_*p*_ represents the vector of standardised joint SNP effects and *ϵ*_*p*_ the vector of normally distributed residuals with variance 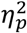. Denote the SNP LD matrix as *S* = *cor(X)* and the vector of estimated marginal SNP effects 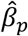 (standardised); we obtain the estimated joint effects from these GWAS summary statistics as 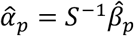, using the reference data set to compute *S*. Here it is assumed that the SNP LD in the reference data is the same as in the original GWAS sample. We can then estimate the residual variance as 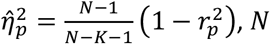 being the original GWAS sample size and *K* the number of SNP principal components (see below), with explained variance 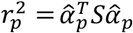. The estimated joint effects 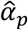 are distributed as 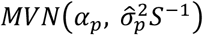, with 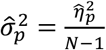 being the sampling variance.

As we cannot be certain whether beta coefficients provided as input are standardised, for each SNP *s*, we create Z-scores using the *p*-value and sign of the provided effect size as 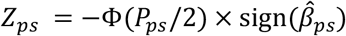, with *Φ* the cumulative normal distribution function and *P*_*ps*_ the SNP *p*-value. We then convert *Z*_*ps*_ to the corresponding correlation 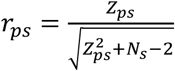, which equals the standardised beta coefficient (note: when per SNP sample size *N*_*s*_ is not provided, we will use the overall *N* as a proxy). If Z-scores or T-statistics are provided we can also use these directly, in which case *p*-values and beta coefficients are not necessary.

Due to the substantial LD between SNPs, it is unlikely that the LD matrix *S* will be of full rank, in which case it is not invertible. This therefore requires us to work in a lower dimensional space. To do so, we compute the singular value decomposition 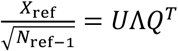, such that *S* = *QΛΛQ*^*T*^ and hence *S*^-1^ = *Q(ΛΛ)*^-1^ *Q*^*T*^ (here *N*_*ref*_ denotes the sample size of the reference data set with genotype matrix *X*_*ref*_). For each component *j*, the corresponding squared singular value 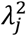 is proportional to the amount of variance of the total accounted for by that component. We order the components by decreasing singular value, and select the smallest subset of the first *K* components such that these account for at least 99% of the total variance (pruning away the rest).

Defining *Q*_∗_ as the *K*_snp_ by *K* pruned eigenvector matrix, and *Λ*_∗_ as the corresponding *K* by *K* diagonal singular value matrix, we approximate the inverse of *S* as 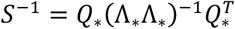. We then define the scaled principal component matrix *W* = *XR* with projection matrix 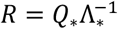. Finally, we define the corresponding vector of joint effects *δ*_*p*_ = *R*^+^*α*_*p*_, with 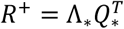, such that *Wδ*_*p*_ closely approximates *G*_*p*_ = *Xα*_*p*_, and use this sparser *δ*_*p*_ in place of *α*_*p*_*/G*_*p*_ for parameter estimation instead.

To test the proportion of phenotypic variance that can be attributed by the local genetic signal, we construct the test statistic 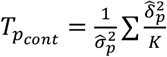 and evaluate this using an F-distribution with *K* and *N* − *K* – 1 degrees of freedom.

### Processing of binary phenotypes

In order to obtain the joint SNP effects from GWAS summary statistics of binary phenotypes, for the scaled principal components *W* as defined above, and with denoting the individual, we reconstruct the multiple logistic regression model 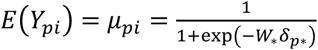 and 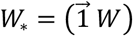, where 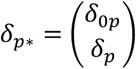, with *δ*_*0p*_ the model intercept. To do so, we use the iteratively reweighted least squares (IRLS) approach^49^, which iteratively updates estimates of the model coefficients according to the equation:

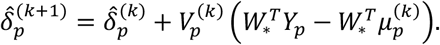

Here, 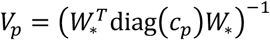, where *c* is a vector with *c*_*pi*_ = _*pi*_ (1 − _*pi*_), and is the index of the current iteration.

For the sufficient statistic 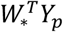 needed for this process, we note that 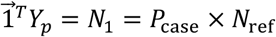 with *P*_case_ designated as the proportion of individuals in the original sample that are cases (*Y*_*pi*_ = 1). In addition, we have that *W*^*T*^*Y*_*p*_ = *R*^*T*^ *X*^*T*^*Y*_*p*_ for the standardised SNP genotype matrix *X* and *R* the projection matrix for *W*. This sufficient statistic can therefore be computed from the individual 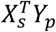 for each SNP *s*. To obtain these, we define the marginal logistic regression model 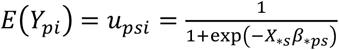, with 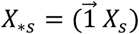 and 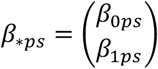, and observe that at convergence of the IRLS algorithm 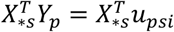 the left side of which can be obtained by filling in the marginal SNP effect estimates 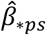. Because the intercept is unlikely to have been reported in the GWAS summary statistics and the slope may not be on the correct scale, we use a search algorithm to re-estimate these SNP effects from the GWAS Z-statistics and case and control counts reported for *s*(substituting general case and control count for the sample if not available per SNP).

From this, the 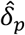 can then be estimated using the IRLS algorithm as outlined above, which has sampling covariance matrix 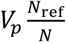. Because the components in *W* are independent and all have the same variance, in practice V_*p*_ should be close to a diagonal matrix, and the standard errors for each 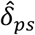 be very similar. To verify this, the ratio between the maximum and median standard error is computed, and if this ratio exceeds 1.75, the PC with the highest standard error is discarded, and the process repeated until no PC has a ratio above that threshold. Subsequently, we define 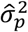 as the mean of the diagonal elements of this sampling covariance matrix, and assume 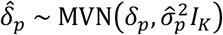 (our simulations show that this approach has no appreciable effect on type 1 error rates, see **Suppl. Fig. 3**). From this we can then also define a test for the univariate signal, similar to the F-test for continuous phenotypes, using the test statistic 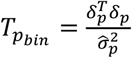. Given the distribution for 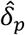, this test statistic has a 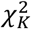 distribution under the null of no genetic association.

### Estimating bivariate local genetic correlations

We define Ω_G_ as the *P* x *P* realised covariance matrix of the genetic components *G* = *Xα* of any *P* phenotypes 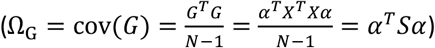, which is the main variable of interest for the estimation of the local genetic correlation. In practice, since we are working with the sparser joint effects of the PCs *δ* (rather than the *α*’ s), which have the same covariance as *G* by a scaling factor *K* (Ω = *α*^*T*^*Sα* = *α*^*T*^*R*^+*T*^*R*^+^*α* = *δ*^*T*^*δ*, and hence 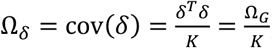), we actually use the Ω_*δ*_ = cov(*δ*) instead. As all the output is standardised, however, this makes no practical difference (since Ω_*G*_ and Ω_*δ*_ have identical correlational structure). We will use Ω to refer to Ω_*δ*_ henceforth. This Ω can be subdivided as:

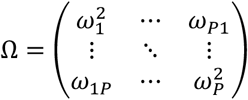

with each diagonal element 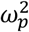 reflecting the (scaled) variance of the genetic component of phenotype *p*, and each off-diagonal element ω_*pq*_ the (scaled) covariance of the genetic components for phenotypes *p* and. We can compute the corresponding bivariate local genetic correlations from the elements of this Ω as 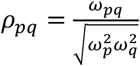 with 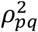 representing the proportion of explained variance (i.e., the local *r*^2^).

For estimation of Ω, we note that the *K* x *P* matrix of estimated joint effects of the principal components 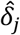 are distributed as 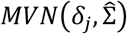 where 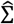 represents the sampling covariance matrix. We then use the Method of Moments^25^ to estimate Ω as follows: With 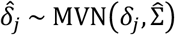, the expected value of 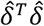 has the form 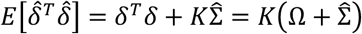, and hence 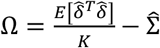. Plugging in the sample moments for 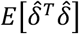, we therefore obtain the estimator 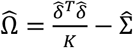.

If there is no sample overlap, 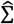 is defined as 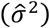, where 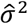 is a length *P* vector with the sampling variances of each phenotype. In the presence of possible sample overlap, estimates of the sampling correlation across phenotypes must be provided by the user. These can be obtained using cross-trait LDSC^13^, creating a *P* x *P* covariance matrix with the intercepts for the genetic covariance for the off-diagonal elements (for the diagonal, use the intercept from a cross-trait analysis of a phenotype with itself, or its univariate LDSC intercept). LAVA then internally converts this to a correlation matrix,, and computes the sampling correlation matrix as 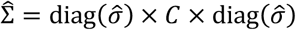.

### Local multiple regression & partial correlations

Local conditional genetic associations between more than two phenotypes can be obtained using either multiple regression or partial correlation.

For the multiple regression approach, consider an outcome phenotype *Y* and set of predictor phenotypes *X*, with corresponding genetic components *G*_*Y*_ and *G*_*X*_. Here, we can decompose *G*_*Y*_ as *G*_*Y*_ = *G*_*X*_ ^*r*)^ into a component that can be explained by *G*_*X*_ and a residual component ε with cov(*G*_*X*_) = 0, such that λ^(r)^ reflects the vector of unstandardised regression coefficients; the variance of is denoted as ε 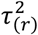. Subdividing 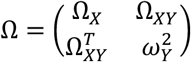 we can then 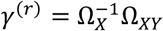 and 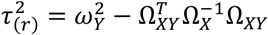 Denoting the vector of standard deviations in Ω_*X*_ as 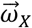, we can then use these to obtain the standardised regression coefficients 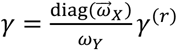 and standardised residual variance 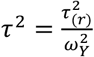. The corresponding explained variance for the full model is computed as *r*^2^ = 1-*τ*^2^.

The partial correlations between the genetic components of two phenotypes *X* and *Y*, conditional on a set of other phenotypes (denoted), can be expressed using the linear equations *G*_*X*_ = *G*_*Z X X*_ and *G*_*Y*_ = *G*_*Z Y Y*_, with _*XY Z*_ = cov(_*X Y*_) As with the parameters from the multiple regression, this can also be computed from the Ω directly. Given the partial covariance 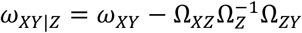 (with subscripts denoting the subset of relevant variances and covariances for *X, Y*, and), and the partial variance 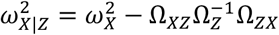 (and likewise for 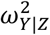), we can simply compute the partial correlation as 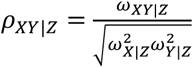.

### Simulation *p*-values and confidence intervals

Because the sampling distributions for the local genetic correlation, partial correlation, and multiple regression coefficients have no tractable closed form, we employ a simulation procedure with partial integration to obtain empirical *p*-values for these parameters. Below, we denote the particular statistic being tested as *T*, with observed value *T*_*obs*_.

First, we define a pure simulation approach, observing that the sufficient statistic 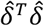 has a noncentral Wishart distribution with *K* degrees of freedom, scale matrix Σ and non-centrality matrix *K*Ω. For a statistic *T*, we can therefore specify the Ω_0_ corresponding to the null hypothesis to be tested and use that to define the non-centrality matrix. We can then generate a random sample of null 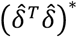 matrices, and for each of those compute the corresponding 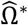 and from there the statistic *T*\*. The sample of null *T*\* values can then be compared to the observed statistic *T*_*obs*_ to obtain an empirical *p*-value, defining this *p*-value as the proportion of simulations for which *T*\* has a value more extreme than *T*_*obs*_.

A drawback of empirical *p*-values is that they can require a substantial number of simulations to reach sufficient accuracy for low *p*-values. To deal with this, we augment the simulation procedure with a partial integration step as follows. For a single phenotype *p*, the distribution of 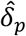 given 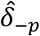 is multivariate normal with parameters of known form, and consequently many of the statistics of interest will have a normal distribution given 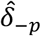 (and Ω_0_). We can therefore generate draws for 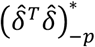 from the noncentral Wishart distribution, and for each such draw compute the parameters of the conditional distribution of the statistic, then obtain the corresponding conditional *p*-value for _*obs*_ for that draw. We then compute the final *p*-value as the mean of the conditional *p*-values across all draws.

Although the resulting *p*-value is still empirical and subject to simulation uncertainty, with this procedure we can obtain sufficiently reliable *p*-values even at very low value ranges without needing prohibitively many simulations. By default, LAVA performs 10,000 simulations to estimate the *p*-value. This is increased this to 100,000 or 1,000,000 simulations if the *p*-value estimate falls below thresholds of 1e-4 and 1e-6 respectively.

For a pair of phenotypes *p* and *q*, to test the null hypothesis of no local correlation, _0 *pq*_ = _*pq*_ =, we use the local covariance _*pq*_ as the statistic to test. For the integration step, to ensure symmetry, we use the conditional distribution of 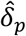 given 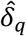 for half of the simulations, and the distribution of 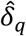 given 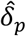 for the other half. Similarly, to test the null hypothesis of no local partial correlation given a set of phenotypes, _0 *pq Z*_ = _*pq Z*_ =, we use the local partial covariance _*pq Z*_ as the statistic *T*. We use the conditional distribution of 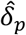 given 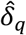 and 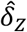 for half the simulations, and the conditional distribution of 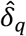 given 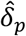 and 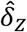 for the other half. For the regression model, with outcome phenotype *Y* and set of predictor phenotypes *X*, to test the null hypothesis of no conditional effect for predictor phenotype j, 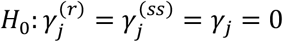, we use the semi-standardised coefficient 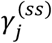 (standardised for *X*, but not *Y*) as the statistic. For the integration step, we use the conditional distribution of 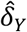 given 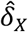.

Optionally, LAVA can also be requested to generate 95% confidence intervals for the local correlation, partial correlation and standardised regression coefficients, as well as for the multiple ^2^ parameter of the multiple regression model. These are computed by generating 10,000 draws from the noncentral Wishart distribution (with non-centrality matrix 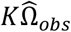), and for statistic of interest computing the simulated statistics *T* * for all draws. The 2.5% and 97.5% quantiles of these *T* * are then used as estimates of the boundaries of the confidence interval for *T*_*obs*_.

### Genome partitioning

In order to partition the genome into smaller regions, we developed a method that uses the LD information between SNPs and groups them into approximately equal sized, semi-independent blocks (available for download at https://github.com/cadeleeuw/lava-partitioning).

The blocking procedure is as follows: For each chromosome, a break point metric for each pair of consecutive SNPs is computed. Starting with the whole chromosome as the initial block, the blocks are then recursively split into two smaller blocks using this metric and a minimum size requirement, continuing until some threshold for the break point metric or size are met and the blocks cannot be divided any further.

Each pair of consecutive SNPs defines a potential break point, for which a metric is computed to determine which breakpoint is the most suitable (i.e., at which point the LD between the SNPs is the lowest). The break point metric between each SNP pair can be thought of as the strength of the LD between the SNPs on each side of the break point. For computational efficiency, we compute only the correlations near the diagonal of the entire SNP x SNP matrix, i.e., between the most proximal SNPs (in this case, we used a window of 200 SNPs). Each break point defines a triangular wedge on this thick diagonal, and the break point metric is simply the mean of the squared correlations in this wedge.

When a block is split, the minimum size requirement is first used to determine the region within the block that contains the subset of potentially valid break points, and within this region, the break point with the lowest metric value is identified. If this value is above some user-defined maximum, the block will not be split any further.

A small margin was also applied to the minimum break point in a block, treating all other potential breakpoints with metrics within that margin as equivalent. The break point closest to the centre of the block was then selected to split the block, in order to encourage more even sizes of sub-blocks. Prior to applying this algorithm, SNPs with a MAF smaller than 1% were filtered out to speed up computation time. These SNPs were added back in after the blocks had been created, applying a variation of the same algorithm to further refine the boundaries between the blocks.

For this paper, we used the default values of the program for all parameters (see program manual for details), except the minimum size requirement which was set to 2500 SNPs in order to obtain an average block size of around 1Mb.

### Simulations and model evaluation

Simulations were conducted in order to validate the robustness of our models, examining the influence of heritability, block size, sample overlap, allele mismatch, and case/control ratio. To ensure an ecologically valid LD structure for our simulations, we used real genotype data from the 1,000 genomes (European subset), from which we simulated phenotypes under various scenarios. In order to achieve a larger sample size than the standard *N* = 503, we stacked the sample 40 times, and subsetted to the first 20,000 individuals. Univariate power for a given locus is fully determined by the sample size and univariate joint effect size (*h*^2^ or OR). Consequently, simulation conditions at N = 20,000 and a particular effect size are representative of conditions at higher N and lower effect size that have the same level of power; for example, for continuous phenotypes with *h*^2^ values of 1%, 5%, 10%, and 25% approximately equivalent power is obtained at an N of 100,000 with *h*^2^ values of .2%, 1%, 2%, and 6% respectively (see **Suppl. Note 3**). For this reason, we opted to keep the sample size constant, and only varied the effect sizes. The original 1,000 genomes sample (*N*=503) was also used as a LD reference for the analysis of the simulated data. Note that as we did not seek to evaluate the influence of mismatching LD between the original data and the reference data, we used the same data set for simulation and analysis in order to prevent unknown violations of model assumptions from influencing the results. Though an extensive investigation of the effects of LD mismatch would be worth undertaking in the future.

The simulations were based on 5 randomly selected loci. Locus size was varied by resizing these 5 loci from the centre SNP and outward until the desired size was achieved (50, 500, 1000, or 5000 SNPs). SNPs with MAF <.01 or an SD of 0 were excluded. All simulations were repeated 1,000 times per block by default, though this was increased to 10,000 for some conditions in order to evaluate type 1 error at lower significance levels.

To evaluate the type 1 error rate for the standard bivariate local *r*_g_ analysis, we simulated two phenotypes with a true local genetic correlation of 0, quantifying the proportion of times where a significant local genetic correlation was detected at different significance levels (*p* < .05, *p* < .01, and *p* < .001). Estimation bias was assessed by simulating true local genetic correlations *ρ* of 0 and .5, and comparing the distribution of estimated correlations to their true value.

For the multiple linear regression, we simulated two genetically correlated (at *ρ* = .5) predictor phenotypes, *X*_0_ and *X*_+_, exhibiting true *r*_g_’s with an outcome phenotype *Y* of 0 and .5, respectively. Detection of a significant effect of *X*_0_ in the multivariate model was considered a false positive, and we used the estimated betas for both *X*_0_ and *X*_+_ to evaluate bias.

For the partial genetic correlations, we generated four predictor phenotypes X, *Y, Z*_1_ and *Z*_2_ simultaneously, with *δ* such that 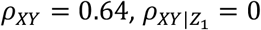, and 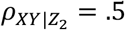. This was accomplished by first generating unit-variance *δ*_*z*1_ and *δ*_*z*2_ such that 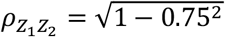 then setting 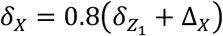 and 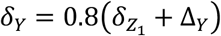 with for the noise terms 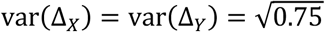 and Δ_*X*_ ⊥ Δ_Y_. Here, we used the *p*-values for 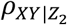 to evaluate the type 1 error rate, and the estimated values for both 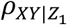 and 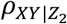 to evaluate bias.

The actual phenotype data was simulated as follows: first, the 1,000 Genomes^24^ genotype data was read in to R (using the snpStats package) and standardised (after increasing of *N* as explained above). We subsequently computed the scaled principal components *W* as defined previously, and for the bivariate and multiple regression simulations, we used these to create the desired Ω (see **Suppl. Note 4** for more detail). With this, we then generated the true *δ*, with which we compute the genetic components *G* = *Wδ*. We obtained the residual variance 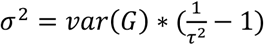, and drew the residuals *ϵ* from a multivariate normal with covariance *σ*^/^*C* (with *C* being the residual correlation matrix used to indicate degree of sample overlap). The *δ* were scaled such that var(*G*) = 1.

For continuous phenotypes, we then generated the *N* × *P* phenotype matrix *Y* as *Y* = β_5_*G* + *ϵ*, with 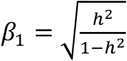 and h^2^ the desired local heritability value. The *P* × 1 vector of residuals *ϵ*_*i*_ for individual *i* was drawn from a normal distribution with zero mean and covariance matrix *C*. The matrix *C* was set to the desired residual correlation matrix for conditions simulating sample overlap, and to *I*_*P*_ otherwise.

For binary phenotypes, the outcome *Y*_*pi*_ for phenotype *p* and individual *i* was modelled as a Bernouilli random variable with probability π_*Pi*_ defined using a logistic model: logit (π_*Pi*_) = β_0_ + β_1_*G*. As for the continuous phenotypes, the β_1_ parameter was used to control the effect size, defining β_1_ = log (*OR*) with *OR* the odds ratio relative to a 1 SD change in the genetic component *G*. The intercept β_0_ was used to control the population prevalence under the model. Its value was determined using a simple linear search, selecting β_0_ such as to obtain the desired prevalence.

### SNP Alignment

As misalignment of SNPs can cause noticeable type 1 error inflation (**Suppl. Fig. 1**), LAVA performs alignment of SNP effect alleles for all summary statistics prior to analysis. This is done first by removing any SNPs with strand ambiguous alleles or alleles that are not present in the reference data set; then, in the case that the reported effect allele does not correspond to that of the reference data set, the sign of the marginal SNP effect size 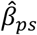 (for phenotype *p* and SNP *s*) is flipped.

### GWAS summary statistics & LD reference data

The GWAS Atlas^2^ (https://atlas.ctglab.nl) was used to search for and access publicly available summary statistics for the 20 traits analysed here, details of which can be found in **Table 1**. We aimed to select a combination of health related and behavioural traits, with the intention of selecting a number of related traits from different categories (e.g., immune, cardiovascular, body composition, psychiatric) in order to facilitate the detection of local genetic correlations, while maintaining some level of phenotypic diversity. When imputation quality metrics were available, we filtered out any SNPs with an INFO score < .9.

As a reference for the estimation of LD in all of our analyses, we used the European subset of the 1,000 Genomes^24^ data as downloaded from https://ctg.cncr.nl/software/magma.

### Global genetic correlation analysis and estimation of sample overlap using LDSC

Bivariate LD-score regression (LDSC)^13^ was used to evaluate the global *r*_g_’s between all trait pairs, as well as to obtain estimates of their level of sample overlap (as required for our LAVA analyses). To account for the sample overlap, we created a (symmetric) matrix based on the intercepts from the bivariate LDSC analyses (the diagonals populated by the intercepts from the analysis of each phenotype with itself). This was then converted to a correlation matrix and provided to LAVA (see ‘Methods: Estimating bivariate local genetic correlations’, for an overview of how LAVA uses this information). Summary statistics for each phenotype were munged using HapMap SNPs.

## Supporting information

Supplementary Tables 1-81 (hotspots with network plots)

## Data availability

All analyses in this study relied on publicly available summary statistics downloaded from the GWAS Atlas^2^ (https://atlas.ctglab.nl; original sources and Atlas-IDs are referenced in **Table 1**). The locus file used for all the LAVA analyses can be downloaded from https://github.com/josefin-werme/LAVA.

## Code availability

The LAVA software is implemented as an R package which is publicly available at https://github.com/josefin-werme/LAVA. The method used for genome partitioning can be downloaded from https://github.com/cadeleeuw/lava-partitioning.

## ACKNOWLEDGEMENTS

This work was funded by COSYN (Comorbidity and Synapse Biology in Clinically Overlapping Psychiatric Disorders: Horizon 2020 Program of the European Union under RIA grant agreement 667301 to D.P.) and the Netherlands Organization for Scientific Research (NWO: VICI 435-14-005). The analyses were carried out on the Genetic Cluster Computer, which is financed by the Netherlands Organization for Scientific Research (NWO: 480-05-003), by the VU University (Amsterdam, The Netherlands) and the Dutch Brain Foundation, hosted by the Dutch National Computing and Networking Services SurfSARA.

## AUTHOR CONTRIBUTIONS

J.W, S.vd.S, D.P, & C.d.L conceived of the study. J.W and C.d.L developed the statistical framework and implemented the software. J.W performed the analyses, simulations, and wrote the manuscript. J.W, S.vd.S, D.P, & C.d.L participated in the interpretation of the results and revision of the manuscript. All authors provided meaningful contributions at each stage of the project.

## References

1. Chesmore, K., Bartlett, J. & Williams, S. M. The ubiquity of pleiotropy in human disease. Hum. Genet. 137, 39–44 (2018).

2. Watanabe, K. et al. A global overview of pleiotropy and genetic architecture in complex traits. Nat. Genet. 51, 1339–1348 (2019).

3. Visscher, P. M. et al. 10 Years of GWAS Discovery: Biology, Function, and Translation. Am. J. Hum. Genet. 101, 5–22 (2017).

4. Shikov, A. E., Skitchenko, R. K., Predeus, A. V. & Barbitoff, Y. A. Phenome-wide functional dissection of pleiotropic effects highlights key molecular pathways for human complex traits. Sci. Rep. 10, 1037 (2020).

5. Hackinger, S. & Zeggini, E. Statistical methods to detect pleiotropy in human complex traits. Open Biology 7, 170125 (2017).

6. Zhao, W. et al. Identification of new susceptibility loci for type 2 diabetes and shared etiological pathways with coronary heart disease. Nat. Genet. 49, 1450–1457 (2017).

7. Giambartolomei, C. et al. A Bayesian framework for multiple trait colocalization from summary association statistics. Bioinformatics 34, 2538–2545 (2018).

8. Hormozdiari, F. et al. Colocalization of GWAS and eQTL Signals Detects Target Genes. Am. J. Hum. Genet. 99, 1245–1260 (2016).

9. Porcu, E. et al. Mendelian randomization integrating GWAS and eQTL data reveals genetic determinants of complex and clinical traits. Nat. Commun. 10, 3300 (2019).

10. Solovieff, N., Cotsapas, C., Lee, P. H., Purcell, S. M. & Smoller, J. W. Pleiotropy in complex traits: Challenges and strategies. Nature Reviews Genetics 14, 483–495 (2013).

11. Pickrell, J. K. et al. Detection and interpretation of shared genetic influences on 42 human traits. Nat. Genet. 48, 709–717 (2016).

12. Giambartolomei, C. et al. Bayesian Test for Colocalisation between Pairs of Genetic Association Studies Using Summary Statistics. PLoS Genet. 10, e1004383 (2014).

13. Bulik-Sullivan, B. et al. An atlas of genetic correlations across human diseases and traits. Nat. Genet. 47, 1236–41 (2015).

14. Zeng, P., Hao, X. & Zhou, X. Pleiotropic mapping and annotation selection in genome-wide association studies with penalized Gaussian mixture models. Bioinformatics 34, 2797–2807 (2018).

15. Grotzinger, A. D. et al. Genomic structural equation modelling provides insights into the multivariate genetic architecture of complex traits. Nat. Hum. Behav. 3, 513–525 (2019).

16. Yang, J., Lee, S. H., Goddard, M. E. & Visscher, P. M. GCTA: A tool for genome-wide complex trait analysis. Am. J. Hum. Genet. 88, 76–82 (2011).

17. van Rheenen, W., Peyrot, W. J., Schork, A. J., Lee, S. H. & Wray, N. R. Genetic correlations of polygenic disease traits: from theory to practice. Nat. Rev. Genet. 20, 567–581 (2019).

18. Zheng, J. et al. LD Hub: A centralized database and web interface to perform LD score regression that maximizes the potential of summary level GWAS data for SNP heritability and genetic correlation analysis. Bioinformatics 33, 272–279 (2017).

19. Shi, H., Mancuso, N., Spendlove, S. & Pasaniuc, B. Local Genetic Correlation Gives Insights into the Shared Genetic Architecture of Complex Traits. Am. J. Hum. Genet. 101, 737–751 (2017).

20. Lu, Q. et al. A Powerful Approach to Estimating Annotation-Stratified Genetic Covariance via GWAS Summary Statistics. Am. J. Hum. Genet. 101, 939–964 (2017).

21. Frei, O. et al. Bivariate causal mixture model quantifies polygenic overlap between complex traits beyond genetic correlation. Nat. Commun. 10, 2417 (2019).

22. Zhang, Y. et al. Local genetic correlation analysis reveals heterogeneous etiologic sharing of complex traits. Preprint at bioRxiv (2020). doi:10.1101/2020.05.08.084475

23. Guo, H., Li, J., Lu, Q. & Hou, L. Detecting Local Genetic Correlations with Scan Statistics. Preprint at bioRxiv (2019). doi:10.1101/808519

24. The 1000 Genomes Project Consortium. A global reference for human genetic variation. Nature 526, 68–74 (2015).

25. Newey, W. K. & West, K. D. Hypothesis Testing with Efficient Method of Moments Estimation. Int. Econ. Rev. (Philadelphia). 28, 777 (1987).

26. Canela-Xandri, O., Rawlik, K. & Tenesa, A. An atlas of genetic associations in UK Biobank. Nat. Genet. 50, 1593–1599 (2018).

27. Schunkert, H. et al. Large-scale association analysis identifies 13 new susceptibility loci for coronary artery disease. Nat. Genet. 43, 333–340 (2011).

28. Liu, J. Z. et al. Association analyses identify 38 susceptibility loci for inflammatory bowel disease and highlight shared genetic risk across populations. Nat. Genet. 47, 979–986 (2015).

29. Schafmayer, C. et al. Genome-wide association analysis of diverticular disease points towards neuromuscular, connective tissue and epithelial pathomechanisms. Gut 68, 854–865 (2019).

30. Pulit, S. L. et al. Meta-Analysis of genome-wide association studies for body fat distribution in 694,649 individuals of European ancestry. Hum. Mol. Genet. 28, 166–174 (2019).

31. Okada, Y. et al. Genetics of rheumatoid arthritis contributes to biology and drug discovery. Nature 506, 376–381 (2014).

32. Julià, A. et al. Genome-wide association study meta-analysis identifies five new loci for systemic lupus erythematosus. Arthritis Res. Ther. 20, 100 (2018).

33. Bronson, P. G. et al. Common variants at PVT1, ATG13-AMBRA1, AHI1 and CLEC16A are associated with selective IgA deficiency. Nat. Genet. 48, 1425–1429 (2016).

34. Howard, D. M. et al. Genome-wide meta-analysis of depression identifies 102 independent variants and highlights the importance of the prefrontal brain regions. Nat. Neurosci. 22, 343–352 (2019).

35. Nagel, M., Watanabe, K., Stringer, S., Posthuma, D. & Van Der Sluis, S. Item-level analyses reveal genetic heterogeneity in neuroticism. Nat. Commun. 9, 905 (2018).

36. Lee, J. J. et al. Gene discovery and polygenic prediction from a genome-wide association study of educational attainment in 1.1 million individuals. Nat. Genet. 50, 1112–1121 (2018).

37. Bulik-Sullivan, B. K. et al. LD Score Regression Distinguishes Confounding from Polygenicity in Genome-Wide Association Studies. Nat. Genet. 47, 291–295 (2015).

38. Flajnik, M. F. & Kasahara, M. Comparative genomics of the MHC: Glimpses into the evolution of the adaptive immune system. Immunity 15, 351–362 (2001).

39. Matzaraki, V., Kumar, V., Wijmenga, C. & Zhernakova, A. The MHC locus and genetic susceptibility to autoimmune and infectious diseases. Genome Biology 18, 76 (2017).

40. Ferreira, M. A. R. et al. Genetic Architectures of Childhood- and Adult-Onset Asthma Are Partly Distinct. Am. J. Hum. Genet. 104, 665–684 (2019).

41. Wen, W. et al. Genome-wide association studies in East Asians identify new loci for waist-hip ratio and waist circumference. Sci. Rep. 6, (2016).

42. Wu, L. et al. Copy number variations of HLA-DRB5 is associated with systemic lupus erythematosus risk in Chinese Han population. Acta Biochim. Biophys. Sin. (Shanghai). 46, 155–160 (2014).

43. Lam, T. H., Tay, M. Z., Wang, B., Xiao, Z. & Ren, E. C. Intrahaplotypic variants differentiate complex linkage disequilibrium within human MHC haplotypes. Sci. Rep. 5, 1–16 (2015).

44. Zhou, H. et al. Genome-wide meta-analysis of problematic alcohol use in 435,563 individuals yields insights into biology and relationships with other traits. Nat. Neurosci. 23, 809–818 (2020).

45. Pasman, J. A. et al. GWAS of lifetime cannabis use reveals new risk loci, genetic overlap with psychiatric traits, and a causal influence of schizophrenia. Nat. Neurosci. 21, 1161–1170 (2018).

46. Nagel, M. et al. Meta-analysis of genome-wide association studies for neuroticism in 449,484 individuals identifies novel genetic loci and pathways. Nat. Genet. 50, 920–927 (2018).

47. Lane, J. M. et al. Genome-wide association analyses of sleep disturbance traits identify new loci and highlight shared genetics with neuropsychiatric and metabolic traits. Nat. Genet. 49, 274–281 (2017).

48. Mota, N. R. et al. NCAM1-TTC12-ANKK1-DRD2 gene cluster and the clinical and genetic heterogeneity of adults with ADHD. Am. J. Med. Genet. Part B Neuropsychiatr. Genet. 168, 433–444 (2015).

49. Agresti, A. An introduction to categorical data analysis. Statistics in Medicine (John Wiley & Sons, Inc, 2007).

