## Supplementary Material for "LAVA: An integrated framework for local genetic correlation analysis"

### SIMULATION RESULTS

#### Overview

This section contains all figures for the simulations results presented in this paper. Given the increased computational burden for simulating and obtaining summary statistics for binary phenotypes, simulations have been conducted using continuous phenotypes unless otherwise specified. By default, locus size has been set to 1000 SNPs, and the number of iterations per locus/condition to 1000 for each of the 5 loci (i.e. 5,000 iterations in total).

### Bivariate model

#### *Allele misalignment*

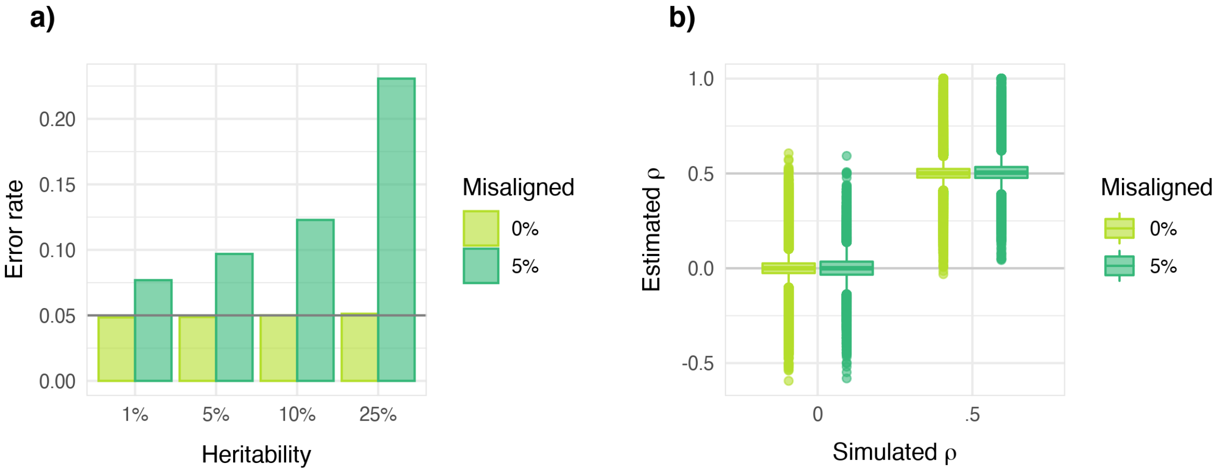

***Supplemental Figure 1. Bias and type 1 error rates as a result of misaligned alleles between reference data and summary statistics**. Type 1 error rates at α = 0.05 for the bivariate local genetic correlation (plot **a**), together with the bias in estimated correlation coefficients ρ (plot **b**), are shown when 0% of alleles have been misaligned, compared to 5%. While misalignment does not lead to bias in the estimated correlation coefficients ρ, it does cause inflation of the type 1 error rate, and for this reason SNP alignment has been implemented as an internal pre-processing step in LAVA.*

#### *Continuous phenotypes*

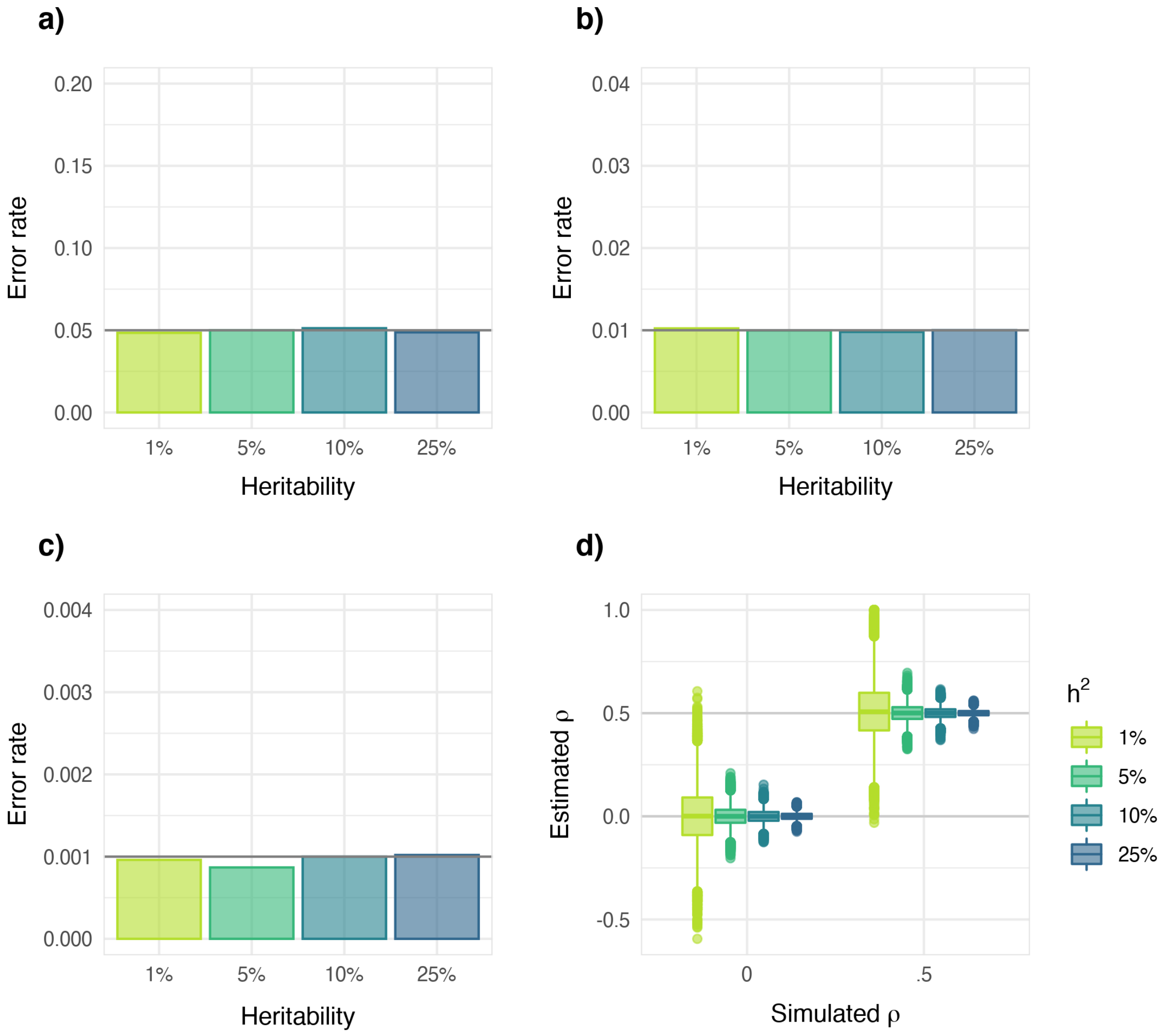

***Supplemental Figure 2. Bias and type 1 error rates for the estimated bivariate genetic correlation across different levels of alpha**. Plot **a-c** shows the type 1 error rates across different levels of heritability for significance threshold $\alpha$ of 0.05, 0.01 and 0.001, respectively (each Y-axis has been scaled to run from 0 to $4\alpha$). Additional iterations have been added to increase resolution at lower alphas (a total of 10,000 per locus, i.e., 50,000 per scenario), and as shown, error rates are contained at the level of $\alpha$ at all thresholds. Plot **d** shows the estimated bivariate correlation coefficients $\rho$ against the simulated $\rho$, indicating that the estimates vary around their true value and are thus unbiased.*

#### *Binary phenotypes*

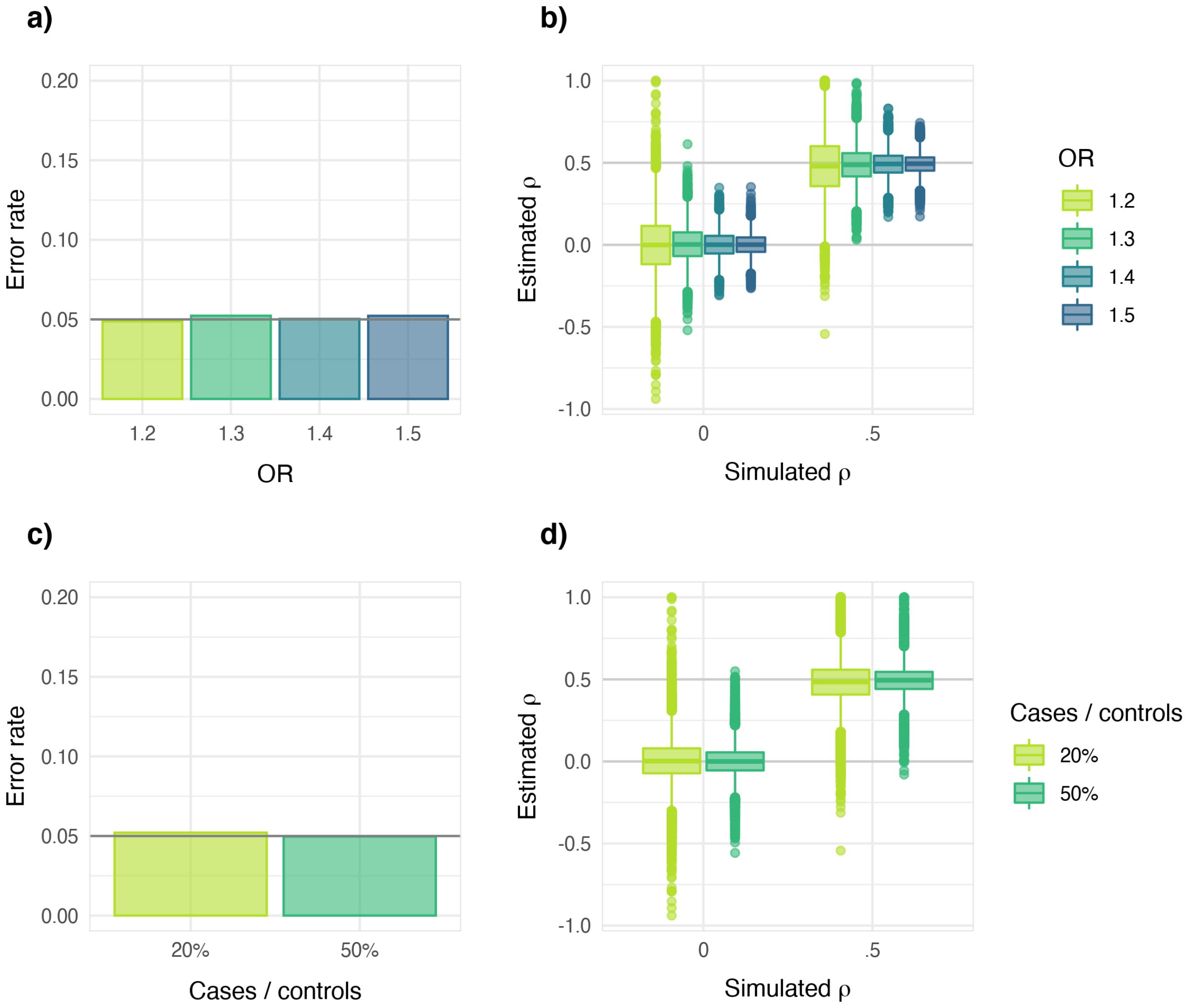

***Supplemental Figure 3. Bias and type 1 error rates for the estimated bivariate genetic correlations with binary phenotypes.*** *Plot **a** and **c** show the type 1 error rates for different ORs and case/control ratios, respectively, at significance threshold $\alpha$ of 0.05 (the Y-axis has been scaled to run from 0 to 4$\alpha$). These plots indicate that the error rates are contained at the level of $\alpha$ across settings. Plot **b** and **d** show the estimated bivariate correlation coefficients $\rho$ against the simulated $\rho$ for different ORs and case/control ratios, indicating that the estimates vary around their true value and are thus unbiased.*

#### *Locus size*

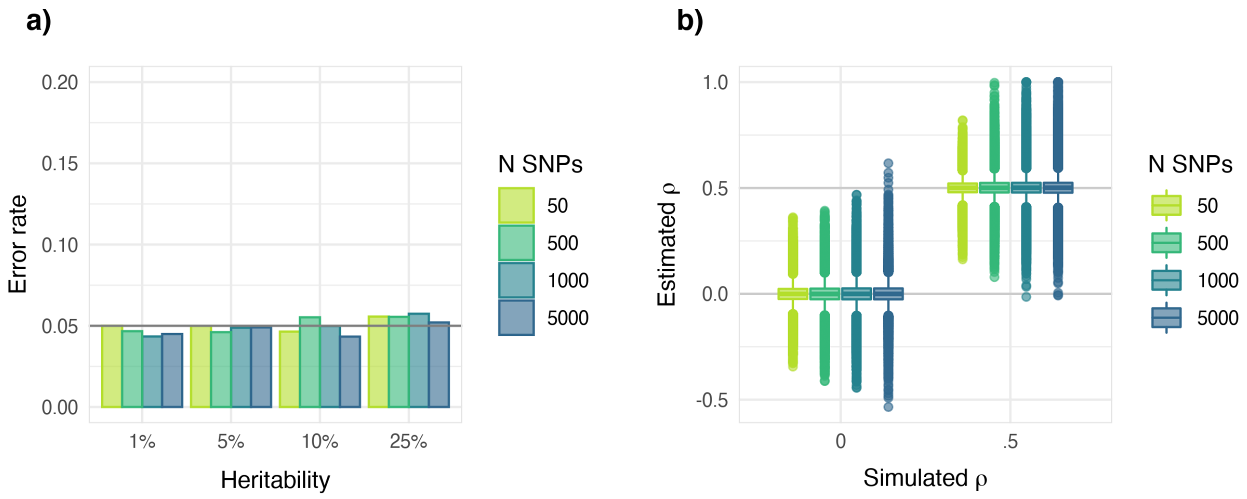

***Supplemental Figure 4. Bias and type 1 error rates for genomic loci of different sizes**. Plot **a** shows the type 1 error rate at α = 0.05 for different levels of heritability for varying locus sizes. Plot **b** shows the estimates of ρ per locus size and heritability. Locus size was varied by taking the centre point of each locus and expanding the boundaries outwards equally on both sides such that the total number of SNPs within the locus arrive at 50, 500, 1000, or 5000 SNPs. As shown, varying the locus size does not lead to bias in the estimated coefficients, or inflated type 1 error rates.*

#### Sample overlap

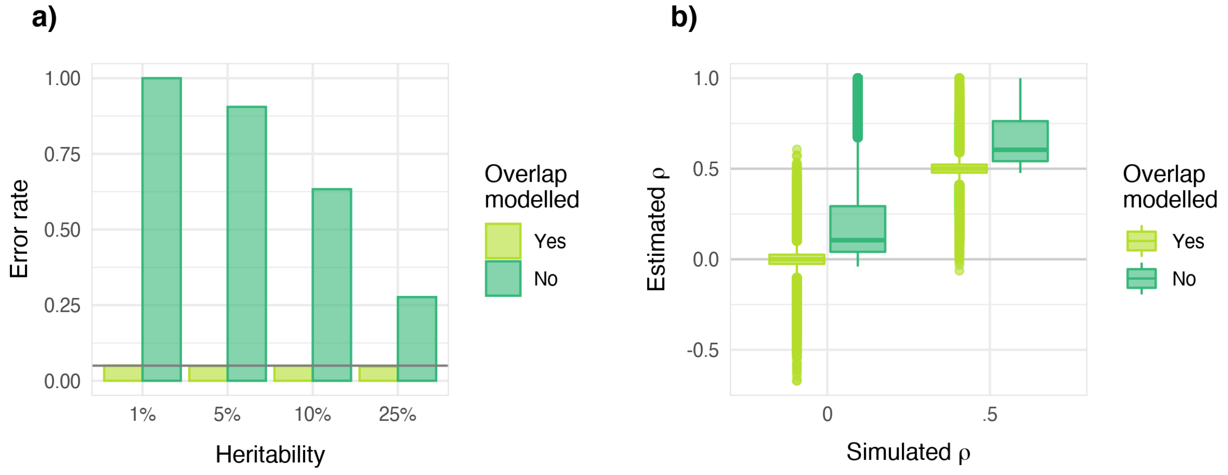

***Supplemental Figure 5. Bias and type 1 error rates for summary statistics with overlapping samples.*** *Sample overlap was simulated by setting the residual phenotypic correlation between data sets to .5. Plot **a** shows the influence of this accounting for or ignoring this sample overlap on the type 1 error rate at $\alpha$ = 0.05; and plot **b** illustrates the effect of accounting for or ignoring sample overlap on the estimated local genetic correlation $\rho$. As shown, not properly accounting for sample overlap leads to both inflated type 1 error, as well as a biased in the estimated coefficients.*

### Multivariate models

#### *Multiple linear regression*

##### Continuous phenotypes

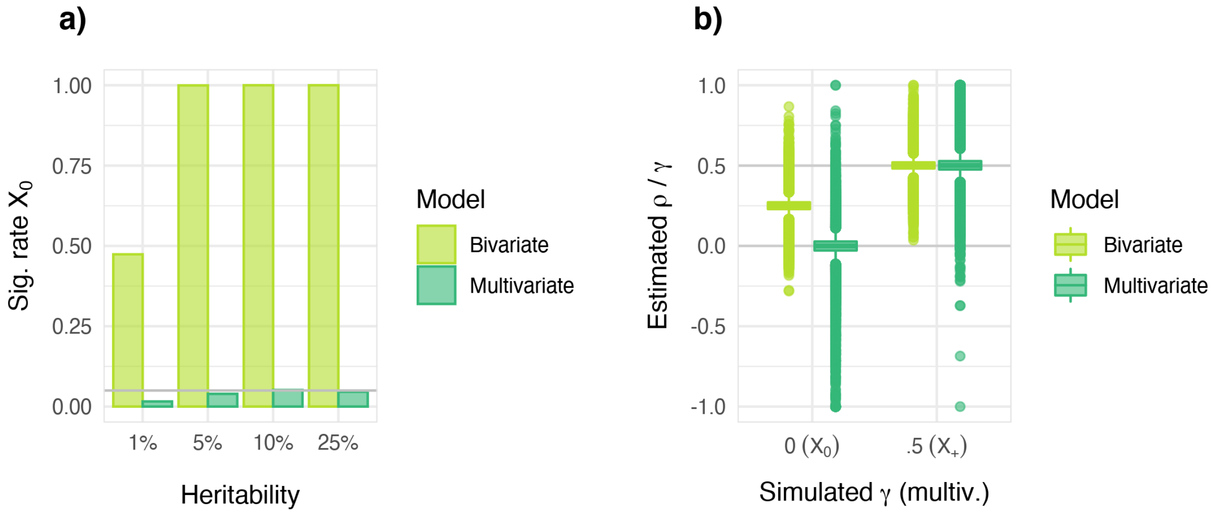

***Supplemental Figure 6. Bias and type 1 error rates for local genetic multiple regression model with two correlated predictors (contrasted against the simple bivariate model) for continuous phenotypes.*** *Here, two predictors phenotypes, $X_0$ and $X_+$, have been simulated with true joint $\gamma$'s of 0 and .5 with the outcome $Y$ (respectively) and a covariance of .5. In plot **a**, the significance rates at $\alpha$ = 0.05 for the relation between $X_0$ and $Y$ under either model, with box-plots of the estimates in plot **b**. As shown, in this setup, the marginal correlation $\rho$ between $X_0$ and $Y$ will be (correctly) estimated as non-zero and significant when using the simple bivariate model (since its relation to the true predictor $X_+$ is unaccounted for). But when analysed jointly together with $X_+$ in the multiple regression model, the conditional association between $X_0$ and $Y$ given $X_+$ is correctly estimated to be 0, with corresponding type 1 error rates equal to $\alpha$. The association between $X_+$ and $Y$ is correctly estimated as 0.5 in both models.*

#### Binary phenotypes

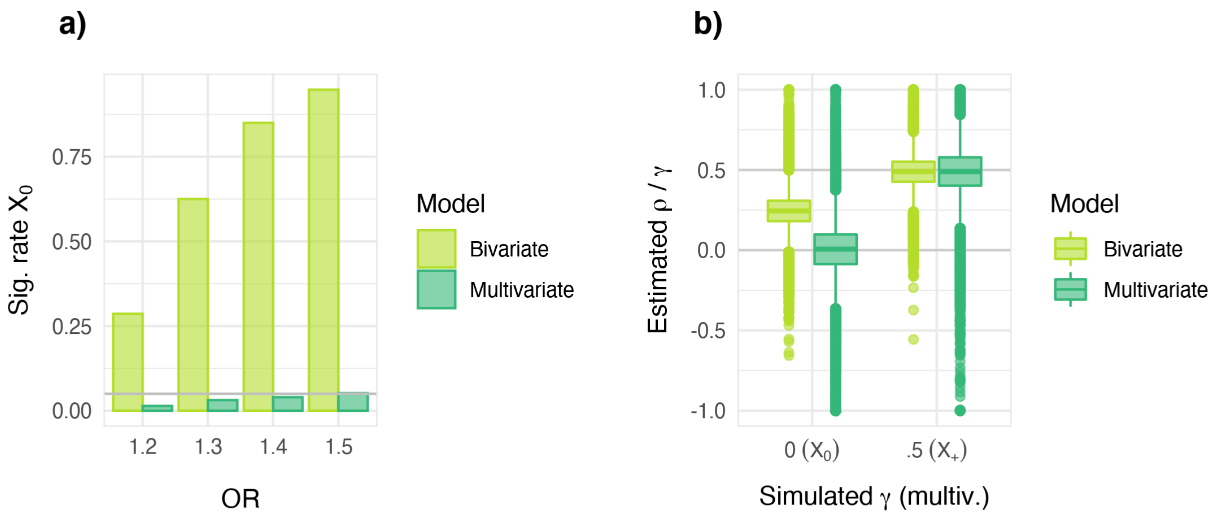

***Supplemental Figure 7. Bias and type 1 error rates for local genetic multiple regression model with two correlated predictors (contrasted against the simple bivariate model) for binary phenotypes.*** *Here, two predictors phenotypes, $X_0$ and $X_+$, have been simulated with true joint $\gamma$'s of 0 and .5 with the outcome $Y$, respectively, and a covariance of .5. In plot* ***a****, t he significance rates at $\alpha$ = 0.05 for the relation between $X_0$ and $Y$ under either model, with box-plots of the estimates in plot* ***b****. As shown, in this setup, the marginal correlation $\rho$ between $X_0$ and $Y$ will be (correctly) estimated as non-zero and significant when using the simple bivariate model (since its relation to the true predictor $X_+$ is unaccounted for). But when analysed jointly together with $X_+$ in the multiple regression model, the conditional association between $X_0$ and $Y$ given $X_+$ is correctly estimated to be 0, with corresponding type 1 error rates equal to $\alpha$. The association between $X_+$ and $Y$ is correctly estimated as 0.5 in both models.*

#### *Partial correlation*

##### Continuous phenotypes

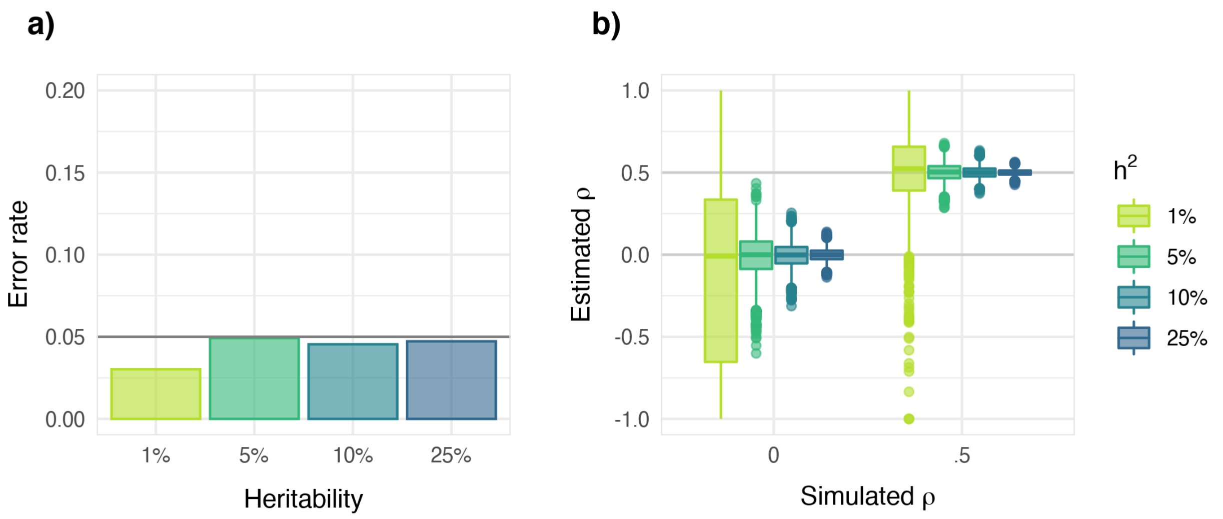

***Supplemental Figure 8. Bias and type 1 error for the partial local genetic correlation using continuous phenotypes.*** *Here, partial correlations have been simulated for two target phenotypes, conditioned on a third. Plot **a** shows the type 1 error rate at α = 0.05 when this partial correlation is simulated at 0, indicating that error rates are contained at α; and plot **b** shows the estimated partial correlation coefficients ρ (compared to the simulated ρ), indicating that the estimates vary around their true value and are thus unbiased.*

#### Binary phenotypes

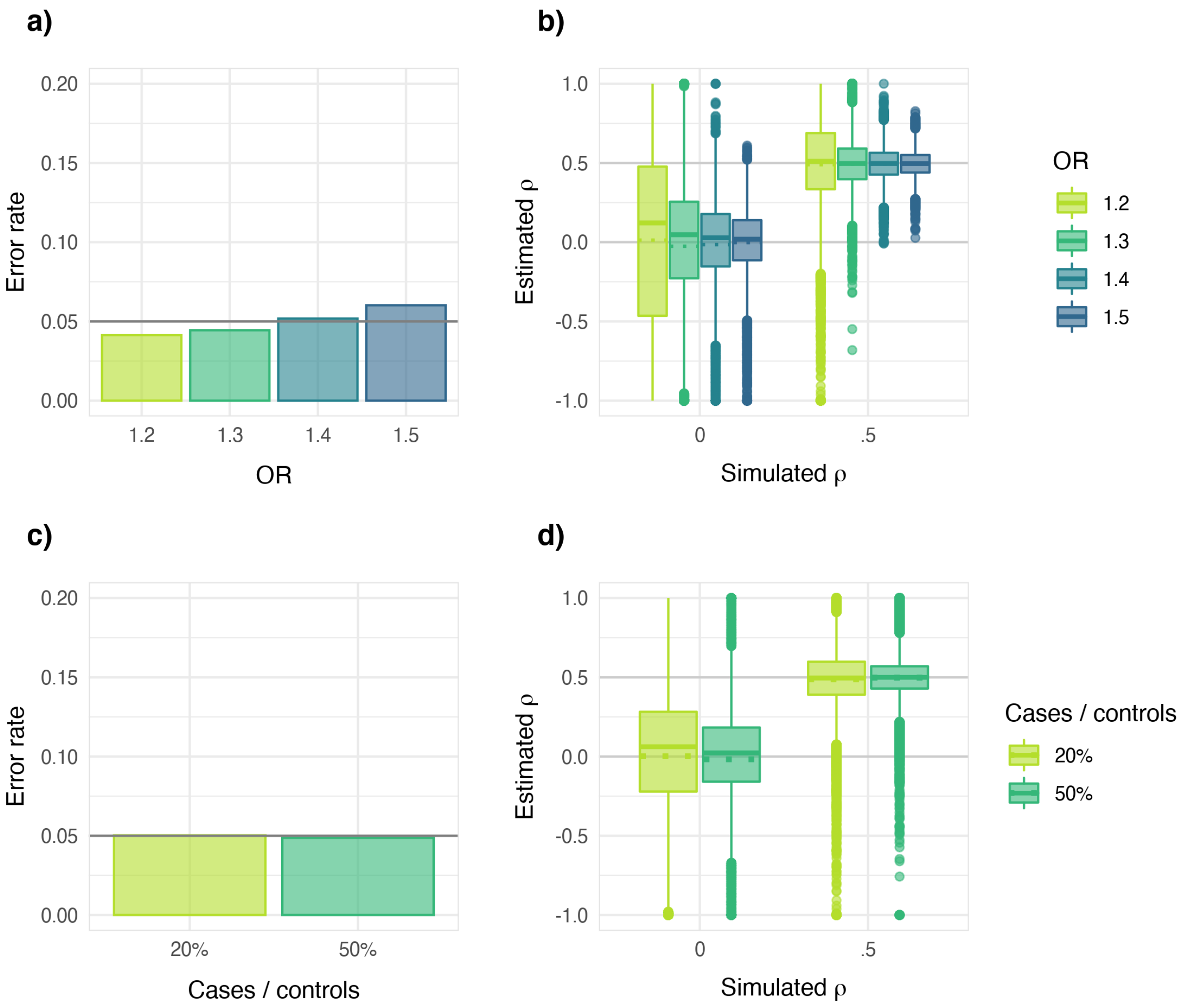

***Supplemental Figure 9. Bias and type 1 error for the partial local genetic correlation using binary phenotypes.*** *Here, partial correlations have been simulated for two target phenotypes, conditioned on a third. Plots **a** and **c** show the type 1 error rate at α = 0.05 when the partial correlation is simulated at 0, indicating that error rates are generally contained at α; the only exception being for OR = 1.5 where there appears to be a very slight inflation (α = .06). We note, however, that this level of effect within a single locus is quite extreme for a complex, non-mendelian traits. Plots **b** and **d** show the estimated partial correlation coefficients ρ, compared to the simulated ρ. Here, we did note a slight bias in the median estimated parameters in the null simulations for lower ORs and case/control ratio*

*($\rho$ = 0; solid lines), though no evidence of a bias in the mean of the estimated parameters (dotted lines) was found, and their type 1 error rates are nevertheless well-controlled.*

### SUPPLEMENTARY RESULTS

#### Direction of significant local $r_g$'s detected with LAVA

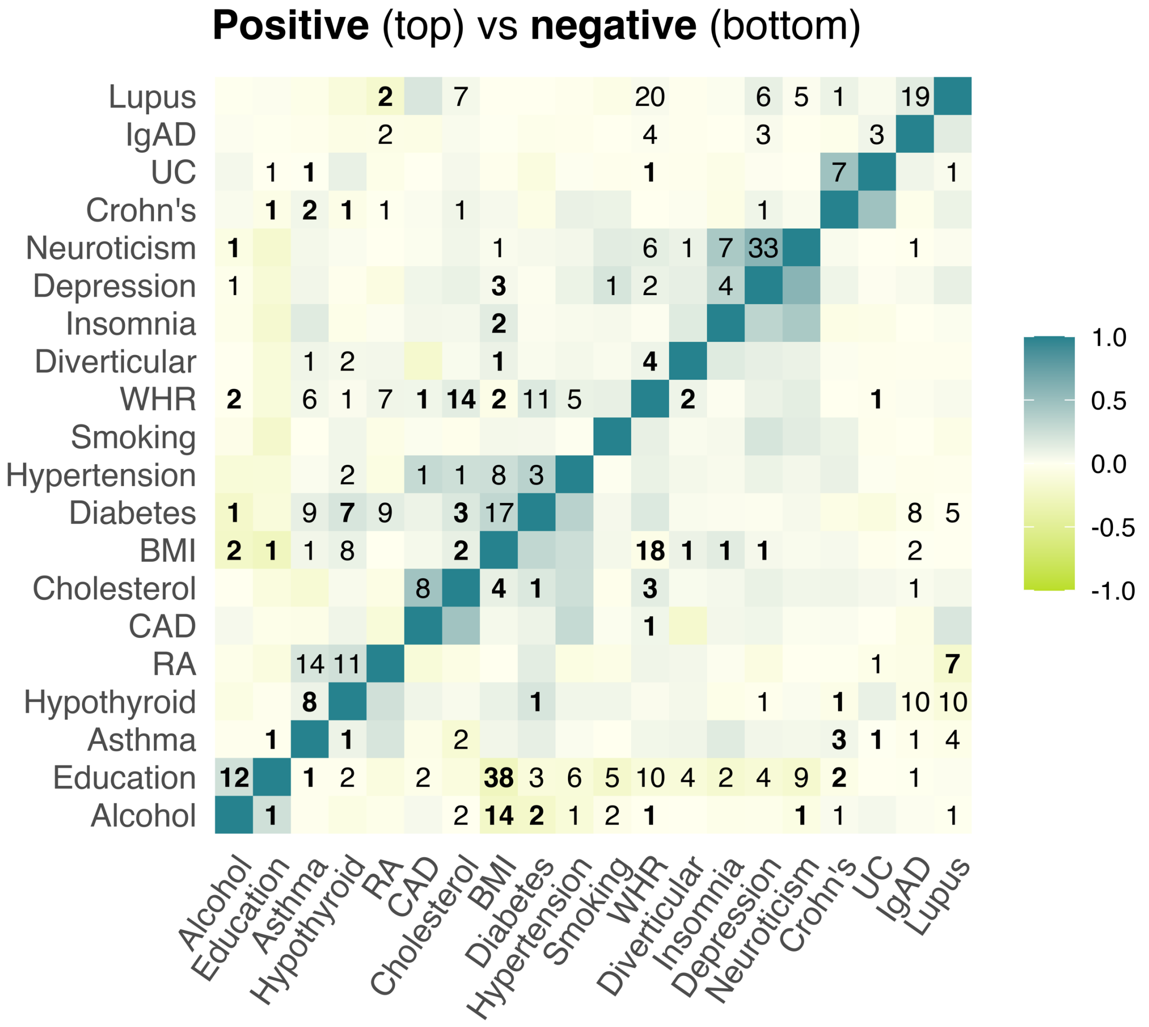

***Supplemental Figure 10. Heatmap showing the number of significant positive (top) and negative (bottom) local genetic correlations detected with LAVA.*** *Values in bold indicate that both positive and negative local $r_g$'s were detected for that trait pair (at $p < .05 / 20{,}630 = 2.42e\text{-}6$), while the colours reflect the mean genetic correlation across tested loci.*

### SUPPLEMENTARY NOTES

#### 1. Comparison of bivariate local $r_g$ estimation used in LAVA, Rho-Hess & SUPERGNOVA

This section contains a general comparison of the bivariate local genetic correlation model in LAVA, with those of rho-HESS[1] and SUPERGNOVA[2] [version published on BiorXiv 10 May 2020].

To facilitate the comparison, we will assume that the data are exactly identical after pre-processing, and whenever possible, equations and formulas from the rho-HESS and SUPERGNOVA papers have been translated to the notation used in this paper. Page and/or equation numbers in the original papers will be referenced for clarity. Note that the comparisons are made only for analysis of continuous phenotypes, as the rho-HESS and SUPERGNOVA models make no specific accommodations for binary phenotypes.

For reference, we will repeat some of the definitions and notation of the LAVA model: For each continuous phenotype $p$ we assume a linear model $Y_p = X\alpha_p + \epsilon_p$, with $X$ a standardised genotype matrix, $Y_p$ the standardised phenotype vector, $\alpha_p$ the vector of joint genetic effects, and $\epsilon_p$ a normally distributed vector of residuals with mean of 0 and variance $\eta_p^2$. We use $\beta_p$ to denote the vector of marginal genetic effects, $N$ and $K_{\text{snp}}$ to denote the sample size and number of SNPs in $X$ respectively, $\eta_{pq}$ for the residual covariance of two phenotypes $p$ and $q$, and $\varphi_{pq}$ for the full covariance between $Y_p$ and $Y_q$.

We define $S = \text{cor}(\text{X})$ as the local LD matrix, which is of rank $K$. We also note the decomposition $Q\Lambda\Lambda Q^T$, with $Q$ the matrix of eigenvectors and $\Lambda$ the diagonal matrix of singular values. We define the standardised principal components $W = XR$ with $R = Q\Lambda^{-1}$ with their genetic effects $\delta$ (such that $W\delta_p = X\alpha_p$). We can now express the relation $\beta_p = S\alpha_p$ and similarly $\alpha_p = S^{-1}\beta_p = R\delta_p$. For the bivariate model, the genetic covariance matrix $\Omega_G = \delta^T\delta$, with estimator

$\widehat{\Omega}_G = \hat{\delta}^T\hat{\delta} - K\hat{\Sigma}$, and $\hat{\Sigma}$ being the sampling covariance matrix which is diagonal in the absence of sample overlap.

#### *Rho-HESS*

In the absence of sample overlap, the estimator for the local genetic covariance in rho-HESS between phenotypes $p$ and $q$ is defined as $\widehat{\omega}_{pq} = \hat{\beta}_p^T S^{-1}\hat{\beta}_q$ (Eq. 5, p. 740). From the equations above, we have $\hat{\beta}_p^T S^{-1}\hat{\beta}_q = \hat{\alpha}_p^T S S^{-1} S\hat{\alpha}_q = \hat{\alpha}_p^T S\hat{\alpha}_q = \hat{\delta}_p^T R^T S R\hat{\delta}_q = \hat{\delta}_p^T \Lambda^{-1} Q^T Q\Lambda\Lambda Q^T Q\Lambda^{-1}\hat{\delta}_q = \hat{\delta}_p^T\hat{\delta}_q$.

This estimator is therefore identical to the off-diagonal element of $\widehat{\Omega}_G$ as estimated by LAVA. Note that in practice, both LAVA and rho-HESS use truncated SVD to deal with rank-deficiency of $S$ and improve stability, replacing $S^{-1}$ with a pseudo-inverse $S^{+}$, though the criteria used to define the truncation differ.

When accounting for sample overlap, rho-HESS subtracts a bias term from $\widehat{\omega}_{pq}$, where this bias term equals $K(\varphi_{pq} - \widehat{\omega}_{pq})\frac{N_{pq}}{N_p N_q}$, with $N_{pq}$ the number of overlapping individuals (Eq. 6-8, p. 740). This differs slightly from the bias correction in LAVA which equals $K\hat{\sigma}_{pq} = K\sqrt{\sigma_p^2\sigma_q^2}C_{pq} = K\sqrt{\frac{\eta_p^2\eta_q^2}{(N_p-1)(N_q-1)}}C_{pq}$, where $C_{pq}$ is an estimate of the sampling correlation.

Like rho-HESS, LAVA recommends obtaining this using bivariate LDSC[3]. However, rather than using just the bivariate LDSC intercept $i_{pq} = \frac{\varphi_{pq}N_{pq}}{\sqrt{N_p N_q}}$, LAVA also incorporates the univariate LDSC intercepts $i_p$ and $i_q$ as the corresponding variance terms to normalise $i_{pq}$ to account for possible influence of population stratification, defining $C_{pq} = \frac{i_{pq}}{\sqrt{i_p i_q}}$.

In the case that $i_p = i_q = 1$, the bias correction term for LAVA reduces to $K\varphi_{pq}\sqrt{\eta_p^2\eta_q^2}\frac{N_{pq}}{N_p N_q}$ (if we assume for simplicity that $(N_p - 1)(N_q - 1) = N_p N_q$). In this case, effectively the difference is that rho-HESS subtracts the estimated genetic covariance $\hat{\omega}_{pq}$ from the total phenotypic covariance $\varphi_{pq}$ to obtain the residual covariance, whereas instead LAVA uses the $\sqrt{\eta_p^2\eta_q^2}$ to rescale $\varphi_{pq}$ to the scale of the residual covariance. In practice, the two different corrections should be very similar.

To obtain local genetic correlations, the covariance estimate $\hat{\omega}_{pq}$ is simply divided by the corresponding variances, which are defined simply as the local genetic covariance of a phenotype with itself. Like the local covariance estimates, these should also be very similar between the two methods.

For testing significance, rho-HESS assumes that the sampling distributions of the local genetic correlation and covariance are normal, using a parametric bootstrap approach to estimate the standard errors for these sampling distributions. By contrast, LAVA uses a simulation approach based on a non-central Wishart distribution to directly generate *p*-values, since during development normal distributions were found to be insufficiently accurate as approximations to the true sampling distributions, particularly for lower *p*-values.

#### *SUPERGNOVA*

In contrast to both LAVA and rho-HESS, SUPERGNOVA assumes the vector of joint genetic effects $\alpha_p$ for a phenotype $p$ to be random rather than fixed, assuming a bivariate normal distribution for the effects $\alpha_j = \begin{pmatrix}\alpha_{pj} \\ \alpha_{qj}\end{pmatrix}$ for SNP $j$, and an assumption of independence of these across SNPs. This bivariate normal distribution has means of zero, and its covariance is the genetic covariance matrix $\Omega_\alpha$ that is to be estimated.

To estimate this $\Omega_\alpha$ SUPERGNOVA uses the same kind of approach as bivariate LDSC, and thus differs from LAVA (and rho-HESS) in both the underlying model assumptions, as well as the parameter estimation. To facilitate the comparison, we will therefore first derive a Method of Moments style estimator of the kind used by LAVA, but for the model assumed by SUPERGNOVA. We can then compare this to the LAVA estimator to isolate the effect of the difference in model assumptions, given the same approach to estimation.

Under the SUPERGNOVA random effects model, we have $\hat{\alpha}_p \sim \mathrm{MVN}\left(0, \omega_{Sp}^2 I_{K_{\mathrm{snp}}} + \hat{\sigma}_p^2 S^{-1}\right)$. Consequently, it follows that $\hat{\delta}_p = \Lambda Q^T \hat{\alpha}_p$ has a multivariate normal distribution with means of 0 and covariance of $\Lambda Q^T\left(\omega_{Sp}^2 I + \hat{\sigma}_p^2 S^{-1}\right) Q\Lambda = \omega_{Sp}^2 \Lambda\Lambda + \hat{\sigma}_p^2 I$, meaning that in this projected space the individual $\hat{\delta}_{pj}$ are independent of each other. For each component $j$, $\hat{\delta}_j = \begin{pmatrix} \hat{\delta}_{pj} \\ \hat{\delta}_{qj} \end{pmatrix} \sim \mathrm{MVN}\left(0, \lambda_j^2 \Omega_\alpha + \hat{\Sigma}\right)$, where $\lambda_j$ is the singular value for that component (and hence $\lambda_j^2$ is its eigenvalue).

Given this distribution $E\left[\hat{\delta}_j \hat{\delta}_j^T\right] = \lambda_j^2 \Omega_\alpha + \hat{\Sigma}$, we can construct a Method of Moments estimator of the form:

$$\widehat{\Omega}_\alpha = \frac{1}{K} \sum_j \frac{1}{\lambda_j^2} (\hat{\delta}_j \hat{\delta}_j^T - \hat{\Sigma})$$

For comparison, the estimator of $\Omega_G$ in LAVA can be rewritten as:

$$\widehat{\Omega}_G = \hat{\delta}^T \hat{\delta} - K\hat{\Sigma} = \sum_j \left(\hat{\delta}_j \hat{\delta}_j^T - \hat{\Sigma}\right).$$

Because the difference in overall scaling cancels out when looking at the genetic correlation, the only substantive difference is the weighting by $\frac{1}{\lambda_j^2}$ in $\widehat{\Omega}_\alpha$.

It should be noted however, that this weighting is not strictly necessary to obtain an estimate of $\Omega_\alpha$. Since $E\left[\hat{\delta}_j \hat{\delta}_j^T - \hat{\Sigma}\right] = \lambda_j^2 \Omega_\alpha$, it follows that under the random effects assumption $E\left[\widehat{\Omega}_G\right] =$

$\Omega_\alpha \sum_j \lambda_j^2$, and therefore $\frac{\widehat{\Omega}_G}{\sum_j \lambda_j^2}$ is an unbiased estimator of $\widehat{\Omega}_\alpha$ as well (albeit not as statistically efficient as $\widehat{\Omega}_\alpha$). The converse is not true, however, and under fixed effect assumptions the two estimators estimate fundamentally different quantities. With $\alpha$ fixed, $\widehat{\Omega}_\alpha$ becomes an estimator of $\frac{\alpha^T\alpha}{K}$, effectively the realised covariance of $\alpha$. Note that this realised covariance of $\alpha$ still only reflects the covariance of the joint rather than the causal SNP effects: even if the realised correlation of $\alpha$ equals 1, it does not follow that causally speaking the same SNPs are involved for the two phenotypes.

The estimator used in SUPERGNOVA itself is defined in terms of the marginal Z-statistics for the SNPs, with $z_{ps} = \frac{\hat{\beta}_{ps}}{\hat{\sigma}_{ps}}$ for phenotype $p$ SNP $s$ and $\hat{\sigma}_{ps}$ the standard error for that SNP (Suppl. Note 1.3, p. 3). These are then transformed using the eigenvectors of the LD matrix, $\tilde{Z}_p = Q^T Z_p$ (Suppl. Note 1.4, p. 7). The model is then defined akin to the bivariate LDSC model, modelling the expected value of the products of the projected Z-scores. For simplicity of comparison, we will assume there is no sample overlap, in which case we obtain (Suppl. Note 1.5, p. 7):

$$E\left[\tilde{z}_{pj}\tilde{z}_{qj}\right] = \frac{\sqrt{N_p N_q}}{K} l_j^2 \omega_{pq}$$

Here, $l_j = \lambda_j^2$, the eigenvalue of principal component $j$.

This equation can essentially be interpreted as a simple linear regression equation, with outcome $\tilde{z}_{pj}\tilde{z}_{qj}$, predictor $\frac{\sqrt{N_p N_q}}{K} l_j^2$ and slope parameter $\omega_{pq}$ (and no intercept), and SUPERGNOVA indeed uses a weighted linear regression approach to estimate $\omega_{pq}$. Weights $q_j^2$ are set to the approximate inverse variance of $E\left[\tilde{z}_{pj}\tilde{z}_{qj}\right]$. This yields the estimator (Suppl. Note, Eq. 10, p. 8):

$$\widehat{\omega}_{pq} = C \sum_j \frac{l_j^2}{q_j^2} \tilde{z}_{pj} \tilde{z}_{qj}$$

with $C = \frac{K}{\sqrt{N_p N_q}} \left( \sum_j \frac{l_j^4}{q_j^2} \right)^{-1}$.

If we now make a simplifying assumption that the $\hat{\sigma}_{ps}$ for all SNPs have the same value $\hat{\sigma}_p$, we can write $Z_p = \frac{\hat{\beta}_p}{\hat{\sigma}_p}$ and hence $\tilde{Z}_p = \frac{1}{\hat{\sigma}_p} Q^T \hat{\beta}_p = \frac{1}{\hat{\sigma}_p} Q^T S \hat{\alpha}_p = \frac{1}{\hat{\sigma}_p} \Lambda\Lambda Q^T \hat{\alpha}_p = \frac{1}{\hat{\sigma}_p} \Lambda\Lambda Q^T Q \Lambda^{-1} \hat{\delta}_p = \frac{1}{\hat{\sigma}_p} \Lambda \hat{\delta}_p$. As such, under this assumption $\tilde{z}_{pj} \tilde{z}_{qj} = \frac{1}{\hat{\sigma}_p \hat{\sigma}_q} \lambda_j^2 \hat{\delta}_{pj} \hat{\delta}_{qj}$. Plugging this into $\widehat{\omega}_{pq}$, we end up with a sum over the terms $\frac{l_j^3}{q_j^2} \hat{\delta}_{pj} \hat{\delta}_{qj}$, subsuming $\frac{1}{\hat{\sigma}_p \hat{\sigma}_q}$ into $C$.

For the weight term $q_j^2$ used, we have $q_j^2 = (D_p l_j^2 + l_j)(D_q l_j^2 + l_j) = D_p D_q l_j^4 + (D_p + D_q) l_j^3 + l_j^2$, where $D_p = \frac{N_p h_p^2}{K}$ and $h_p^2$ the local heritability (and same for $q$). Consequently, the total weights in the summed terms are

$$\frac{l_j^3}{q_j^2} = \frac{1}{D_p D_q} \frac{1}{\lambda_j^2 + H_j}$$

with $H_j = \frac{D_p + D_q + \frac{1}{\lambda_j^2}}{D_p D_q}$. With this, we then end up with an adjusted estimator

$$\widehat{\omega}_{pq}^* = C^* \sum_j \frac{1}{\lambda_j^2 + H_j} \hat{\delta}_{pj} \hat{\delta}_{qj}$$

where $C^* = \frac{C}{D_p D_q}$.

Although not the same, we can see that this is very similar to the $\widehat{\Omega}_\alpha$ estimator presented above. In practice the term $H_j$ should be relatively small as well; eg. if $D_p = D_q = D$, $H_j = \frac{2}{D} + \frac{1}{D^2 \lambda_j^2}$.

The value of $D$ will tend to be considerably larger than 1, since the product of sample size and local heritability $N_p h_p^2$ generally needs to be rather larger than the number of principal components $K$ for there to be any detectable genetic signal in the locus (and absent strong enough signal, the genetic covariance / correlation is not reliable anyway). $H_j$ could in principle also get quite large if the eigenvalue $\lambda_j^2$ is very small but, like LAVA and rho-HESS, SUPERGNOVA also prunes away components with low eigenvalues prior to analysis, so this should not be an issue.

Note as well, however, that the $C$ and $C^*$ terms will differ somewhat for the covariance and the corresponding two variances, and as such will not entirely cancel out in the correlation estimate.

From the structure of $\widehat{\omega}_{pq}^*$, although by no means identical to $\widehat{\Omega}_\alpha$, we can conclude that it operates on mathematically similar principles and should tend to yield similar estimates. It follows, however, that there may be considerable differences in the local genetic correlation estimates (and hence also their p-values) between LAVA and rho-HESS on the one hand and SUPERGNOVA on the other, depending on how the unique and overlapping genetic signal of the two phenotypes is distributed over the principal components in the locus. This goes hand in hand with the fact that, the distinction between fixed and random effect models aside, the two types of local genetic correlation that are estimated are fundamentally different metrics with different interpretations, that therefore cannot be directly compared. The question of which metric is more suitable for what kind of research question requires further study to find an adequate answer.

#### 2. Partial correlations and multiple regression, overview

This note contains an overview of partial correlation and multiple regression, and how they relate. In general, the partial correlation between two (standardised) variables $X$ and $Y$ given a set of (standardised) variables $Z$ can be defined in terms of two linear regression models. Writing $X = Z\alpha_X + \varepsilon_X$ and $Y = Z\alpha_Y + \varepsilon_Y$, the partial correlation $\rho_{XY|Z}$ is the correlation of the two residual terms, $\rho_{XY|Z} = \mathrm{cor}(\varepsilon_X, \varepsilon_Y)$. As such, the partial correlation reflects correlation between $X$ and $Y$ that cannot be accounted for by $Z$, and a partial correlation of zero therefore implies that the variables in $Z$ jointly capture all of the processes responsible for the correlation between $X$ and $Y$.

We can compare the partial correlation to the coefficient $\beta_X$ in the multiple linear regression model $Y = X\beta_X + Z\beta_{ZY} + \delta_Y$, or similarly $\beta_Y$ in the model $X = Y\beta_Y + Z\beta_{ZX} + \delta_X$. Here, if $X$, $Y$ and $Z$ are all standardised, $\beta_X$ and $\beta_Y$ are standardised regression coefficients, and like the partial correlation $\rho_{XY|Z}$ these reflect a measure of the remaining dependency between $X$ and $Y$ when controlling for the variables in $Z$. Unlike the partial correlation, however, multiple regression is not symmetrical: $\beta_X$ and $\beta_Y$ will generally not have the same value (though their p-values will be identical, and standardised $\beta_X$ and $\beta_Y$ will usually be similar).

The differences between the two can be better understood by considering multiple regression with a residualised phenotype. With outcome $Y$, instead of fitting the regression parameters all at once, we can also first regress the effects of $Z$ out of $Y$ and compute the residualised outcome $Y_{\mathrm{res}} = Y - Z\beta^*_{ZY} = \varepsilon_Y$, then perform the regression $Y_{\mathrm{res}} = X\gamma_X + \delta_Y$. If $X$ and $Z$ are correlated, this essentially attributes all shared effects of $X$ and $Z$ on $Y$ to only $Z$, instead of proportionally distributing it over both (note also that $\gamma_X / SD(\varepsilon_Y)$ yields the semi-partial correlation $\mathrm{cor}(X, \varepsilon_Y)$).

The resulting model can be written as $Y = X\gamma_X + Z\beta^*_{ZY} + \delta_Y$, analogous to the original regression. In both cases, the analysis creates a partitioning of the variance of outcome $Y$, with the regression coefficients more or less reflecting the relative size of the component that is attributed to each predictor variable. Thus, the only difference between the normal multiple regression model and its residualised counterpart is the priority assigned to each predictor in making that partitioning.

Conceptually, the main difference between using regression and partial correlation therefore lies in the fact that regression is inherently asymmetrical, assigning a special role to the outcome $Y$ as the variable to be partitioned and understood. If the roles of $X$ and $Y$ are swapped, this yields a different partitioning that is designed to answer a different research question. Partial correlation, by contrast, is entirely symmetrical, with the quantity of interest being the relation between two variables rather than the effect one has on the other.

Practically, their values can also be very different: $\rho_{XY|Z} = \frac{\mathrm{cov}(\varepsilon_X,\varepsilon_Y)}{SD(\varepsilon_X)SD(\varepsilon_Y)}$, whereas $\gamma_X = \frac{\mathrm{cov}(\varepsilon_X,\varepsilon_Y)}{SD(X)SD(\varepsilon_Y)} = \frac{\mathrm{cov}(\varepsilon_X,\varepsilon_Y)}{SD(\varepsilon_Y)} = SD(\varepsilon_X)\rho_{XY|Z}$. Because $X$ is standardised, $SD(\varepsilon_X)$ reflects the proportion of the variance in $X$ that cannot be explained by $Z$. The partial correlation $\rho_{XY|Z}$ will therefore by definition be higher than $\gamma_X$, a difference that increases the greater the correlation between $X$ and $Z$. This difference reflects the fact that partial correlation essentially estimates the (average) relation between $X$ and $Y$ within each subsets of the data defined by $Z$, whereas regression estimates their relation across those subsets, merely subtracting out the part of $Y$ that $Z$ can account for.

In a LAVA analysis, the choice of multiple regression versus partial correlation thus hinges on the research question it is meant to answer. If the primary interest is in the genetic component of a particular phenotype and the aim is to determine to what extent this can be explained and decomposed in terms of the genetic components of other phenotypes, the multiple regression analysis is most suitable. If instead the interest is generally in the genetic correlations between multiple phenotypes and the degree to which these can be accounted for by other phenotypes, a partial correlation analysis would be preferred. Depending on the study, a combination of both approaches may be also be warranted to investigate different aspects of the same genetic relations.

#### 3. Scaling of power by $h^2$ and $N$

For a standardised continuous phenotype $Y = X\beta + \varepsilon$ for standardised predictor $X$ and error term $\varepsilon$ with variance $\sigma^2$, the estimate $\hat{\beta} \sim \mathrm{N}\left(\beta, \frac{\sigma^2}{N}\right)$. Consequently, $Z = \frac{\hat{\beta}}{\sqrt{\frac{\sigma^2}{N}}}$ has a distribution $\mathrm{N}\left(\frac{\beta}{\sqrt{\frac{\sigma^2}{N}}}, 1\right)$. Moreover, the true explained variance $r^2 = \beta^2$ and the residual variance $\sigma^2 = 1 - r^2$. Denoting the expected value of $Z$ as $\mu$, we then have $\mu^2 = \left(\frac{\beta}{\sqrt{\frac{\sigma^2}{N}}}\right)^2 = \frac{r^2}{1-r^2}N$.

With $\sigma^2$ known, power for the test of $\hat{\beta}$ is therefore fully determined by the parameter $\mu$ (since this defines the whole distribution of the Z-statistics), and hence scenarios with equal $\mu$ will also have equal power. We can therefore use the expression for $\mu$ to compute how $r^2$ should be adjusted to maintain the same level of power given a change in $N$.

In particular, $r^2 = \frac{\mu^2}{\mu^2+N}$. Filling this in for different $h^2 = r^2_{20K}$ values under the simulation $N$ of 20,000, the corresponding $r^2_{100K}$ values that yield the same level of power for e.g. an $N$ of 100,000 are:

$r^2_{20K} = 0.01$, then $r^2_{100K} = 0.002$

$r^2_{20K} = 0.05$, then $r^2_{100K} = 0.0104$

$r^2_{20K} = 0.10$, then $r^2_{100K} = 0.0217$

$r^2_{20K} = 0.25$, then $r^2_{100K} = 0.0625$

### 4. Generating the true $\boldsymbol{\delta}$ from $\boldsymbol{\Omega}$ in simulations

#### *Process genotype data*

1. Read in and standardise genotype data
2. Decompose $X = Q\Lambda Q^T$ and define $Q_*$ as the subset of eigenvalues that explain at least 99% of the variance, with the corresponding $\Lambda_*$. Define $W = XQ_*\Lambda_*^{0.5}$ (i.e., the pruned and scaled principal components).

#### *Create deltas*

For a linear regression of an outcome $Y$ on a set of predictors $X$, with $\Omega = \begin{pmatrix} \Omega_X & \Omega_{XY} \\ \Omega_{XY}^T & \omega_Y^2 \end{pmatrix}$, if we specify $Y$ in terms of the regression coefficient $\gamma$ and residual variance $\tau$, we can write this as $\Omega = \begin{pmatrix} \Omega_X & \Omega_X\gamma \\ \gamma^T\Omega_X & \tau^2 + \Omega_{XY}^T\Omega_X^{-1}\Omega_{XY} \end{pmatrix}$. To obtain the desired $\delta$ matrix from this $\Omega$, we proceed as follows:

1. Create the desired $\Omega_X$ (e.g. a diagonal matrix, if there is no genetic correlation between the predictors in $X$, or with some covariance otherwise). For simplicity, we defined this matrix on a standardised scale to ensure the specified covariances within $X$ were equal to their correlation.
2. Set $\omega_Y = 1$ so that the specified $\gamma$ reflects the exact correlation desired, generate $\Omega_{XY}^T$ and $\Omega_{XY}$, and then create the full $\Omega$ as shown above.
3. Generate a matrix $D$ of the size of $\delta$ and center such that its means are zero.
4. Decompose $\frac{D^TD}{K} = Q_1\Lambda_1Q_1^T$ and compute $\delta_* = DQ_1\Lambda_1^{-0.5}$, which will result in $cov(\delta_*) = I_P$
5. Decompose $\Omega = Q_2\Lambda_2Q_2^T$ and, finally, compute $\delta = \delta_*\Lambda_2^{0.5}Q_2^T$, which will result in a matrix $\delta$ with the $cov(\delta) = \Omega$.

### REFERENCES

1. Shi, H., Mancuso, N., Spendlove, S. & Pasaniuc, B. Local Genetic Correlation Gives Insights into the Shared Genetic Architecture of Complex Traits. *Am. J. Hum. Genet.* **101**, 737–751 (2017).
2. Zhang, Y. *et al. Local genetic correlation analysis reveals heterogeneous etiologic sharing of complex traits*. *Preprint at bioRxiv* (2020). doi:10.1101/2020.05.08.084475
3. Bulik-Sullivan, B. *et al.* An atlas of genetic correlations across human diseases and traits. *Nat. Genet.* **47**, 1236–41 (2015).
